# Corticothalamic Neurons in Motor Cortex Have a Permissive Role in Motor Execution

**DOI:** 10.1101/2022.09.20.508799

**Authors:** Lina Marcela Carmona, Anders Nelson, Lin T. Tun, An Kim, Rani Shiao, Michael D. Kissner, Vilas Menon, Rui M. Costa

## Abstract

The primary motor cortex (M1) is a central hub for motor learning and execution. M1 is composed of heterogeneous cell types, many exhibiting varying relationships to movement. Here, we employed an unbiased screen to tag active neurons at different stages of performance of a motor task. We characterized the relative cell type composition of active neurons across training and identified one cell type consistently enriched as training progressed: corticothalamic neurons (M1^CT^). Using two-photon calcium imaging, we found that M1^CT^ activity is largely suppressed during movement, and this negative correlation with movement scales with movement vigor and augments with training. Closed-loop optogenetic manipulation of this population revealed that increasing M1^CT^ activity during forelimb movement significantly hinders execution, an effect that became stronger with training. Similar optogenetic manipulations, however, had little effect on locomotion. In contrast to M1^CT^ neurons, we observed that M1 corticospinal neurons positively correlate with movement, and that this positive correlation increases with learning. Finally, by examining the connectivity between M1^CT^ and corticospinal neurons, we uncovered that M1^CT^ neurons can suppress M1 corticospinal activity via feedforward inhibition, and that this inhibition scales with training. These results identify a novel permissive role of corticothalamic neurons in movement execution through suppression of inhibition of corticospinal neurons.

## INTRODUCTION

The ability to learn and execute new movements is critical for survival. Precise execution of movements is often acquired via repetition and refinement, a process that depends on activity coordinated throughout the brain and spinal cord^1,2^. Within these circuits, the primary motor cortex (M1) projects directly to the spinal cord, but also to other motor areas in the basal ganglia, thalamus and brainstem, affording it unique control over motor action^3,4^. M1 is engaged during motor learning as well as during the execution of voluntary movements in rodents, non-human primates, and humans^5–12^. Changes in the formation, turnover, and spatial distribution of spines^13–17^, the restructuring of dendritic arbors^18–21^, as well as shifts in the patterns of electrical activity in M1 all affirm an active and critical role of M1 in learning and performance of various motor tasks^11,15,22–27^.

However, M1 cells are transcriptionally and functionally heterogeneous^28–30^, and different cell types exhibit varying relationships to movement during learning and/or execution. Furthermore, many of these populations are intricately connected, generating local networks that could contribute to the overall activity and function of M1. Some recent studies have tried to isolate the role of different M1 cell types in movement or learning by using known cell type markers or projection patterns^22,23,31^. We reasoned that a more unbiased and comprehensive screen of which cell types are engaged during movement, at different stages of training and proficiency, could complement the candidate approach taken to date. We designed an experimental approach leveraging a calcium-dependent photoswitchable indicator in combination with single-cell RNA sequencing to determine which M1 cell types are differentially enriched at different stages of motor training and proficiency. This approach identified several neuron types within M1 with differential activity early versus late in training and led us to uncover that a particular cell type – the M1 corticothalamic neurons – have an unexpected but critical role in movement.

## RESULTS

### M1^CT^ neurons are enriched at late stages of training

We utilized an M1-dependent head-fixed motor task^32^ to screen for cell type engagement at different stages of training. The task consists of forelimb-driven pulls of a small rotary wheel wherein mice had to cross an infrared (IR) beam at the apex of the wheel and make contact with the rungs to commence a trial (Figure 1A). Mice were rewarded when pulls executed within 200ms of trial start reached a velocity threshold. Within two weeks, mice showed an increase in the number of successful trials, as demonstrated by significantly increased percent success over time (p=0.0104, one-way ANOVA, time) (Figure 1B), as well as increased velocity of trials, as indicated by an increase in the maximum velocity reached during all trials in each training session (p=0.0182, one-way ANOVA, time) (Figure 1C,D). For subsequent experiments, we chose day 4 and day 12 as our early and late training time points, respectively (p=0.0336, day 4 v. day 12, one-way ANOVA).

**Figure 1.**
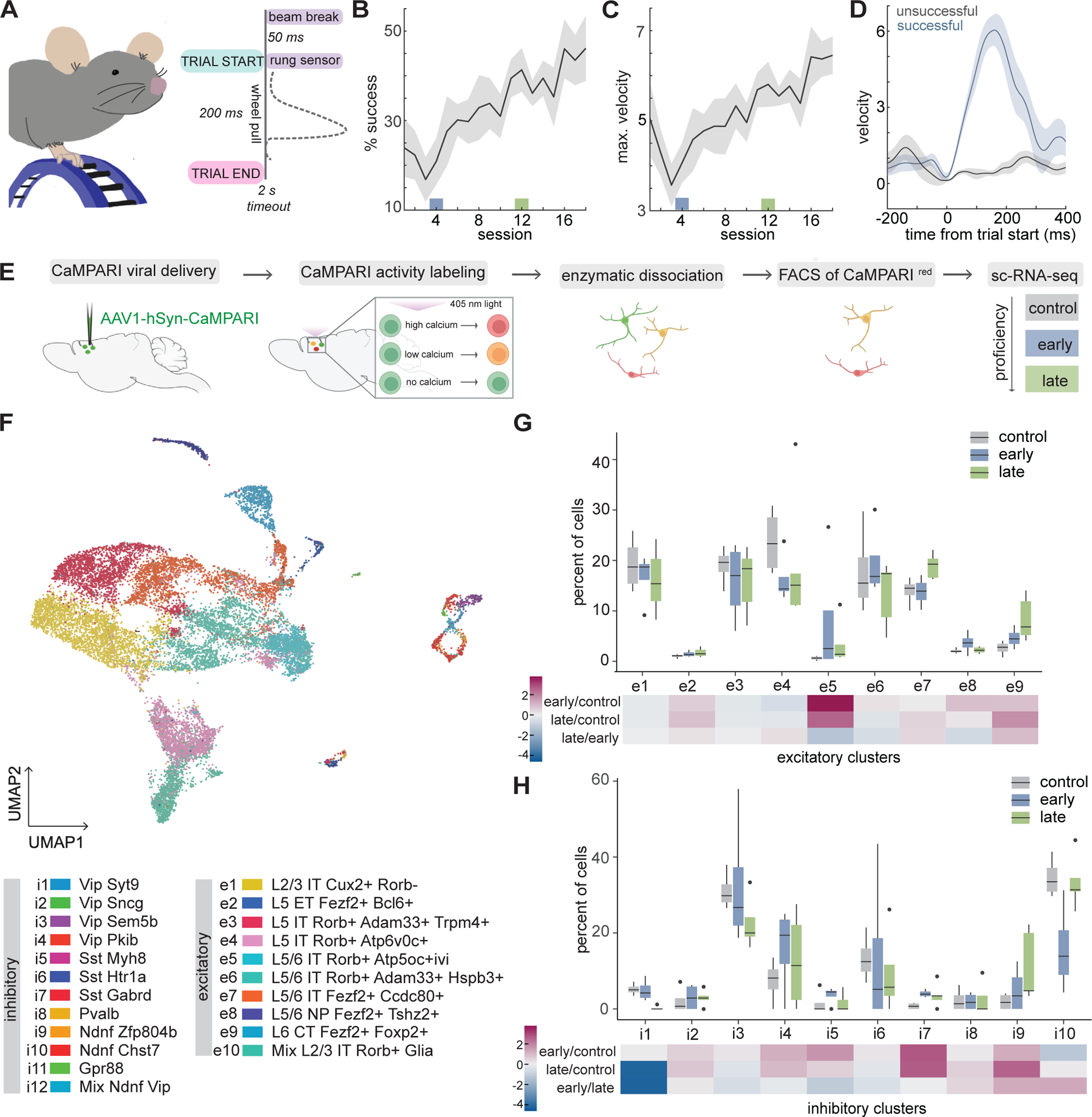
M1 active cell type enrichment during training of a motor task. **A** Schematic of wheel pulling task, left. Schematic of trial structure, right. **B,C** Percent success **(B)** and maximum velocity of all trials **(C)** during 18 consecutive training sessions; N=6 animals. Early and late training are denoted by blue and green boxes, respectively; p=0.0104 **(B)** and p=0.0182 **(C)**, one way ANOVA, time; p=0.0336 **(B)** and p=0.0569 **(C)** early v. late training, one-way ANOVA. **D** Velocity of successful (blue) and unsuccessful trials (gray) on day 4 (early training). **E** Outline of labeling, sorting, and sc-RNAseq strategy. **F** Aggregate UMAP of all sc-RNA-seq runs. **G,H** Percent of excitatory **(G)** or inhibitory **(H)** neurons present from each clusters at each timepoint, top; Gray, control; Blue, early training; Green, late training; outliers, black dots. Fold change between the early and control, late and control, and early and late, bottom.

We next optimized a system to label the neurons that were active at different stages of execution of this task. To this end, we utilized CaMPARI, a photoswitchable indicator that fluoresces green at baseline but converts to red upon coincident exposure to calcium and 405 nm light^33^ (Figure 1E). Although there are numerous systems now available to label active neurons^34–37^, CaMPARI was particularly well suited for our approach given its high fidelity reporting of activity relative to Immediate Early Gene (IEG) systems, and its short window of labeling (on the order of minutes) compared to hours for other systems. We performed stereotactic injections of an adeno-associated virus (AAV) expressing CaMPARI under the neuron-specific human synapsin promoter into the caudal forelimb region^38,39^ of M1. We compared cell type engagement at different training stages: 1) early stages of training (day 4), 2) late stages of training (day 12), and 3) no training, in which the wheel was covered but the mouse was otherwise exposed to the same environment. At the respective time point, light was delivered through a cranial window in pulses of 5s to photoswitch CaMPARI and label the active neurons as mice performed or were exposed to the task (Figure 1S). M1 was then rapidly dissociated and neurons with photoswitched red CaMPARI fluorescence were enriched using fluorescence activated cell sorting (FACS). This ensured that only active neurons were subsequently processed using single-cell RNA sequencing. Each timepoint (control, early, late) was repeated a minimum of four times to account for behavioral variability, with each run consisting of neurons from 2-4 animals. We achieved a median unique molecular identifier (UMI) and gene count of 7971.5 and 3503, respectively, over all runs (Supplementary Table 1).

Unbiased clustering of our aggregate data yielded the expected cell type subdivisions based on cortical layer marker genes. Mapping to a high resolution M1 cell type atlas^28,29^ further demonstrated the cell type heterogeneity present in our samples, with concordance among our clusters and published annotations (Figure 1F, S2A,B). Three clusters, i11, i12, and e10, contained mixed populations and were excluded from further analysis. Comparison across each timepoint did not reveal a binary presence or absence of particular cell types (Figure S2C). Rather, enrichment analysis demonstrated a diversity of cell type engagement patterns across timepoints (Figure 1G,H). Because we did not capture as many inhibitory neurons across timepoints, we conducted our enrichment analysis independently for excitatory and inhibitory neurons. To control for labeling dependency on light penetration and differential baseline activity of neuronal types, we used internal comparisons by calculating enrichment relative to the control condition that did not receive any training. Overall, compared to the control condition that did not receive any training, the most enrichment occurred across clusters in late condition with four subtypes each enriched in excitatory and inhibitory neurons, respectively. In comparison, only two inhibitory and one excitatory subtype were enriched in the early condition relative to the control condition. When comparing early and late training stages, we observed enrichment of three subtypes, one excitatory and two inhibitory, at late stages of training, and enrichment of three inhibitory subtypes at early stages (significance determined using ANCOM-BC pvalue<0.05; Supplementary Table 2). One subtype of particular interest with significant enrichment at the late training timepoint relative to the control (p= 0.003943091) because of its lack of characterization during motor learning and execution was the cluster of corticothalamic neurons (M1^CT^). This population is marked by FoxP2 and Fezf2 expression (excitatory cluster 9, e9) and is experimentally accessible with the FoxP2-cre mouse line. Anatomical mapping of the projections of FoxP2+ M1 neurons confirmed broad thalamic targeting of this population^4,40–42^, and also demonstrated topographical organization in M1 dictated by thalamic projection pattern (Figure S3). Given this enrichment at late stages of learning, as well as the known role of thalamus in motor learning and execution^43–45^, we further investigated the role of these neurons during motor execution.

### M1^CT^ neurons decrease activity during movement

The contribution of the corticothalamic M1 output pathway to motor execution is poorly understood compared to the thalamocortical pathway^46,47^ as well as other output populations in M1^23,27,31^. Thus, we characterized the activity of M1^CT^ neurons at different stages of training in our wheel pulling task. We performed stereotactic injections to deliver an AAV expressing Cre-dependent GCaMP7f to the caudal forelimb area in M1 of FoxP2-cre mice (Figure S4A,B). To image activity of these deep cortical neurons without incurring significant damage to M1, we followed an approach previously implemented to image corticospinal neurons in M1^23,27^. We used two-photon (2p) imaging to record calcium dynamics in the dendritc trunks of FoxP2+ M1^CT^ neurons through a cranial window at least 300 µm below pia. An additional retrograde AAV expressing a Cre dependent red fluorescent protein was injected into the cervical segments of the spinal cord to label the small population of Foxp2 expressing corticospinal neurons and exclude them from our imaging fields (Figure S4A,B). To extract the calcium signal, we used constrained nonnegative matrix factorization (CNMF)^48^, and we excluded any highly correlated units (ρ>0.8) to avoid overrepresentation from branches of the same neurons^23,49^.

We analyzed M1^CT^ neuronal activity during movement by aligning to trial start during successful or unsuccessful trials across all training sessions. Surprisingly, we observed that activity of M1^CT^ decreased during wheel movement (Figure 2A,B; Figure S4C). To examine whether this was a general feature of these neurons, we calculated the cross-correlation of the activity (Δ F/F) to the wheel velocity as measured by the rotary encoder. This further demonstrated a negative relationship between activity and wheel movement throughout the session, with the suppression in neuronal activity slightly preceding peaks in wheel velocity (Figure 2C). Given this striking signature in M1 where other populations exhibit positive correlations with movement^23,27,31^, we examined whether the suppression in M1^CT^ neurons was observed for the majority of M1^CT^ neurons. For this, we examined activity during all wheel pulls of all sessions as we found a similar decrease in activity during movement as in trials (Figure S4D). ROC based classification revealed that most neurons show decreased activity during wheel pulls (69%, M1^CT-down^) relative to a window of low movement prior to the pull. A smaller fraction displayed no modulation (15%, M1^CT-no^ ^mod.^) or increased activity (16%, M1^CT-up^) during wheel movement (Figure 2D-G). To further validate this finding, we performed similar imaging using another well-established marker line for labeling layer VI corticothalamic neurons, Ntsr1-cre (Figure S4E,F). ROC classification corroborated that the majority of M1^CT^ neurons decreased in activity during wheel pulls (Figure S4G).

**Figure 2.**
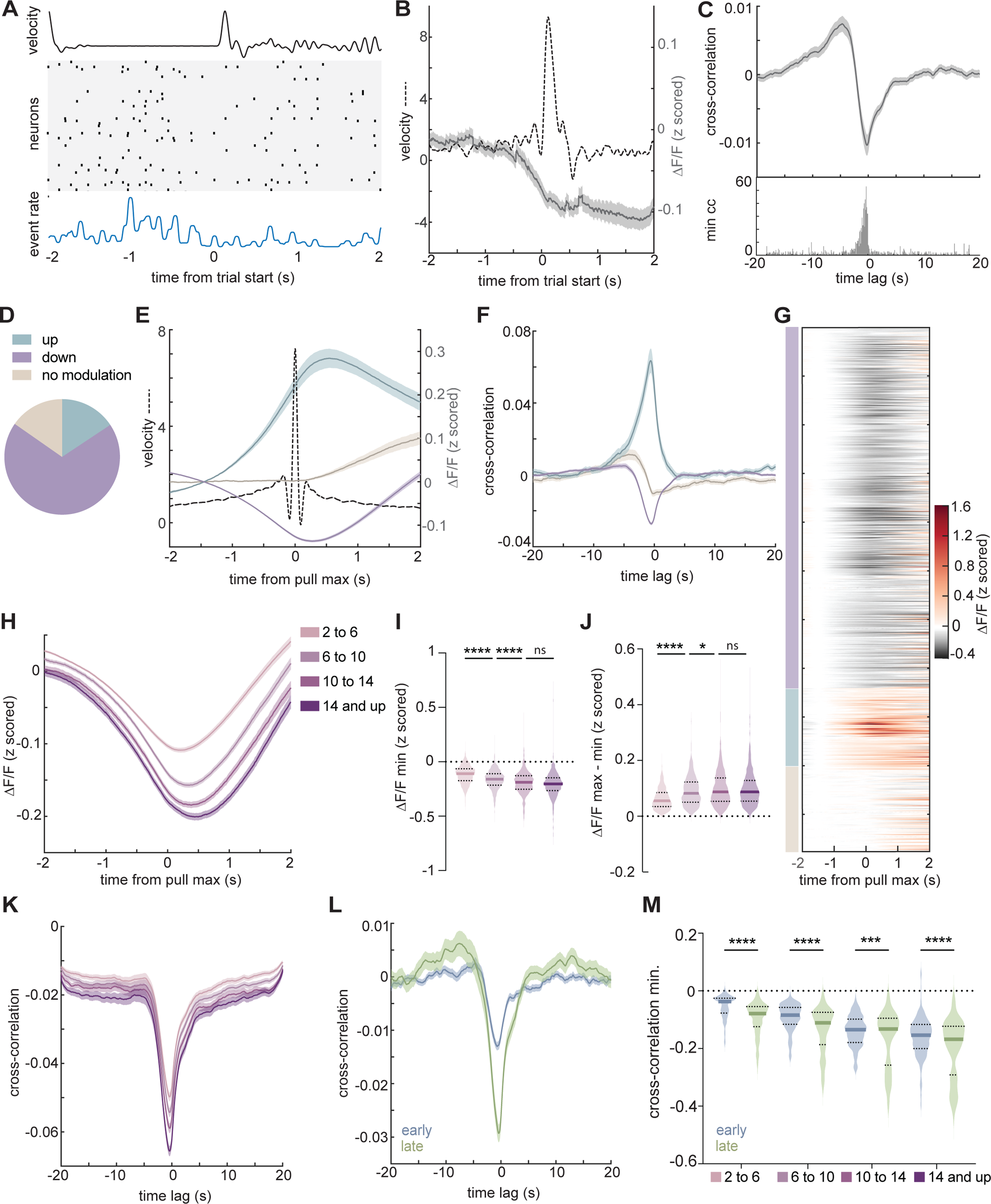
M1^CT^ neurons are suppressed during movement. **A** Single trial example of M1^CT^ activity; top, wheel velocity; middle; raster plot of all units (N=53); bottom, aggregate event rate; 0=trial start. **B** Z-scored ΔF/F during successful trials over all sessions; mean, solid gray line; shaded area, SEM; Wheel velocity during same trials, dotted line; n=1078; N=4; **C** Top, cross-correlation of Z-scored calcium ΔF/F for each unit to wheel velocity for whole session; mean, solid gray line; shaded area, SEM; bottom, binned cross-correlation minimum values at corresponding lags; 50ms bins. **D.** ROC based classification of all units across sessions based on activity during wheel pulls. **E,F** Z-scored ΔF/F **(E)** and cross-correlation to wheel velocity for whole session **(F)** of each group during wheel pulls; solid lines, mean; shaded areas, SEM; wheel velocity during pulls, dotted line in **E**. 0=pull velocity max. **G** Z-scored ΔF/F of all units averaged over all wheel pulls; groups denoted on left. **H** Z-scored ΔF/F of wheel pulls of increasing velocities for down modulated neurons; solid line, mean; shaded area, SEM. **I,J** Distribution of z-scored ΔF/F minimum during wheel pulls **(I)** or the difference in z-scored ΔF/F maximum and minimum before and during wheel pulls **(J)** of down modulated neurons; colors as in **H**. Thick solid line, mean; thin lines, quartiles. One-way ANOVA, multiple comparisons; p<0.0001, 2 to 6 v. 6 to 10 and 6 to 10 v 10 to 14; p=0.0553, 10 to 14 v 14 and up **(I)**, p<0.0001, 2 to 6 v 6 to 10; p=0.230, 6 to 10 v 10 to 14; p=0.9937, 10 to 14 v 14 and up **(J)**. **K,L** cross-correlation to wheel velocity for trials **(K)** and whole session **(L)** for wheel pulls of each velocity bin group **(K)** or early (blue) and late (green) training sessions **(L)** in down modulated neurons; solid line, mean; shaded area, SEM. **M** Distribution of minimum cross-correlation to wheel velocity during pulls of increasing velocity in early (blue) or late (green) sessions of down modulated neurons; thick, line, mean; thin lines, quartiles. Early v. late comparisons, one-way ANOVA, multiple comparisons, p<0.001, 2 to 6, 6 to 10, and 14 and up; p=0.005, 10 to 14. For **B,C**, n=1078 units from N=4 mice. For **D-G**, n=740 downmodulated units, n=168 upmodulated units, and n=168 non-modulated units, all from N=4 mice. For **L,M,** early: n=370 units, late: n= 310 units from N=4 mice.

Furthermore, for the neurons that decreased in activity, the magnitude of the decrease changed depending on the velocity of the movement. When we grouped pulls of varying velocity into bins (2-6 cm/s, 6-10 cm/s, 10-14 cm/s, >14 cm/s), we observed larger decreases in activity for higher velocity wheel movements (Figure 2H-K; for min of ΔF, p<0.0001, 2 to 6 v. 6 to 10 and 6 to 10 v 10 to 14; p=0.0553, 10 to 14 v 14 and up, two-way ANOVA; Figure S5B-C), but no change in the timing of maximal suppression (Figure S5A).

Interestingly, we observed an increase in the modulation range of neurons around movement from early to late training (Δ F/F max – min), consistent with the enrichment in CaMPARI labeling with training (Figure S6A). We therefore examined if the correlation also changed over the span of training sessions and observed a significantly larger anti-correlation during late training sessions compared to early (Figure 2L,M) in M1^CT-down^ neurons. No difference was observed in the relative proportions of the population (Figure S6B). Given that as mice progressed through the task, they tended to perform higher velocity pulls (Figure 1C, Figure S6C), we examined the anti-correlation in early and late session neurons for pulls of matched velocities. Even in subsets of pulls with a similar distribution of peak velocities (Figure S6C), we found a consistent increase in the magnitude of the anti-correlation to wheel movement at late training (Figure 2M, Figure S6D; for early v. late, p<0.001, 2 to 6, 6 to 10, and 14 and up; p=0.005, 10 to 14, one-way ANOVA). These results indicate that M1^CT^ neurons are predominantly suppressed during movement and show a negative correlation with movement that is modulated by speed and increases with training.

### M1^CT^ suppression is permissive for execution of learned movement

We hypothesized that the decrease in M1^CT^ neuronal activity was critical for movement execution during training, and that disrupting this decrease in activity would affect movement. To test this, we performed a closed-loop optogenetic manipulation on a subset of trials at the late training stage where we identified these neurons as most enriched. Given that M1^CT^ neurons decrease in activity during wheel pulls, we chose to activate them during this time window to disrupt this expected decrease. To achieve this, we performed stereotactic injections of an AAV into the caudal forelimb area of M1 to express Cre-dependent channelrhodopsin (hChR2) or a fluorescent protein in Foxp2-cre animals, generating our opsin and control groups respectively (Figure S7A). We then trained both groups of animals to the late training stage and delivered 400 ms of pulsed light (20 hz; 10 ms pulse width) at the onset of one third of trials (Figure 3A). Overall, this manipulation decreased the number of successful trials performed with the light on in opsin expressing animals but not in control animals (Figure 3B,C; for percent of successful trials, p=0.0119, two-tailed unpaired t-test; for percent success, p=0.7031 and p=0.0014 for control and hChR2, respectively, two-way ANOVA). Importantly, when we compared the way in which trials were executed with or without light, we found a drastic decrease in wheel velocity (p=0.8713 and p=0.0003 for control and hChR2, respectively, two-way ANOVA) as well as pull distance (p=0.9511 and p=0.0027 for control and hChR2, respectively, two-way ANOVA) in light-on trials of opsin animals (Figure 3D-F and Figure S7B-G). Furthermore, this decrease in velocity was rapid, and observed in less than 200ms (Figure 3D; p<0.0001 and p>0.9999 for opsin and control, respectively; two-way ANOVA for 400ms after light on, time x condition).

**Figure 3.**
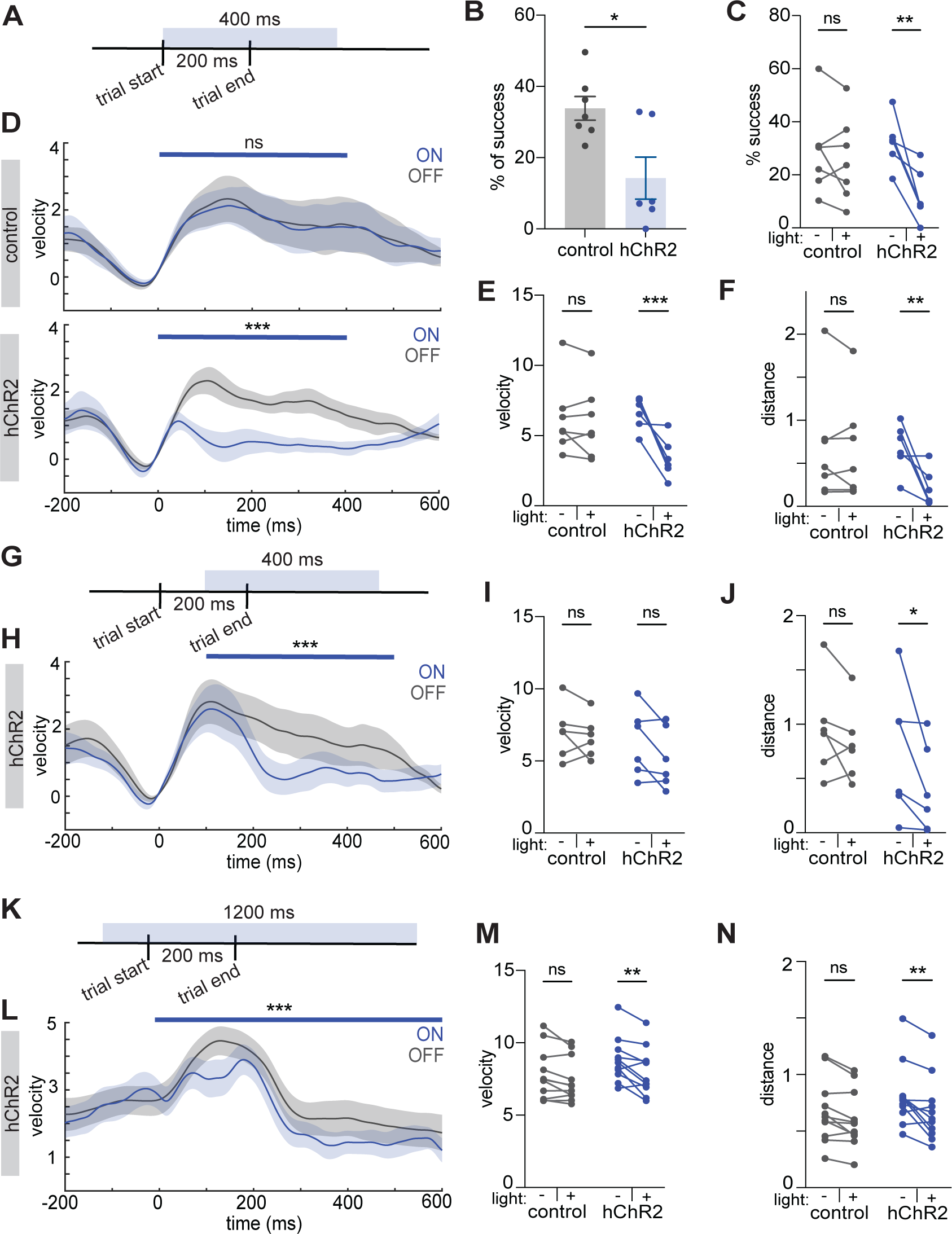
M1^CT^ activity disruption perturbs movement execution. **A, G, K** Schematic of optogenetic light delivery for closed loop at trial start (**A**), closed loop at velocity max (**G**), and open loop trials with light on 100 ms preceding pull start (**K**) for wheel turning task. Light was pulsed at 20 hz; 10 ms pulse width. **B** Percent of successful trials with light on (# successful trials during light on / # of all light on trials); control, gray; hChR2, blue; error bars, SEM; two-tailed unpaired t-test, p=0.0119. For B-J: N=7, control group; N=6, hChR2 group. Each point is average of all light delivery sessions. **C** Percentage of trials that are successful for each animal with light on or off in each group (# (un)successful trials during light on / # of all (un)successful trials); error bars, SEM; two-way ANOVA, multiple comparisons; p=0.7031, control; p=0.0014, hChR2**. D** Velocity traces of all trials from control, top, and hChR2, bottom, animals during light, blue, and no light trials, gray; 0=trial start; blue bar denotes time of light, p>0.9999 (control), p<0.0001 (hChR2), time x condition, two way ANOVA; solid line, mean; shaded area, SEM. **E, I, M.** Maximum velocity of all trials for each animal with light on or off in each group; two-way ANOVA, multiple comparisons. **E**, p=0.8713 (control), p=0.0003 (hChR2)**; I**, p=0.7329, (control), p=0.0789 (hChR2); **M**, p=0.1558 (control, p=0.0016 (hChR2); N=6 for both groups for H-J. N=11 for both groups for L-N **F,J,N** Wheel distance traveled during all trials for each animal with light on or off for each group; two way ANOVA, multiple comparisons; **F**, p=0.9511, (control), p=0.0027, hChR2**; J**, p=0.5128 (control), p=0.0406 (hChR2); **N**, p=0.0708 (control), p=0.0011 (hChR2). **H,L** Velocity traces of all trials from hChR2 animals during light, blue, and no light trials, gray; 0=trial start; blue bar denotes time of light; p=0.0003 **(H)** and p<0.0001 **(L)**, time x condition, two way ANOVA; solid line, mean; shaded area, SEM. See Table 3 for p-values for all other comparisons.

While these results showed that disruption of the activity of these neurons impaired proficient execution of task-related movements, they did not indicate whether modulation of M1^CT^ neuronal activity is important for ongoing movement after peak velocity has been achieved. We, therefore, performed a closed-loop manipulation on a new cohort of animals, delivering light around the time of maximum trial velocity, rather than trial initiation (Figure 3G, light delivered 110 ms after trial start). As expected, there was no change in the percent of successful trials or maximum velocity of trials as mice had already reached peak velocity before the light was turned on (Figure 3I and Figure S8B,C). However, we observed that even after movement had started, there was a rapid and drastic decrease in movement speed following stimulation (Figure 3H,J and Figure S8A,D-I). Finally, we tested whether stimulating these neurons before movement initiation (i.e. to prevent the decrease in M1^CT^ neural activity before movement start) would affect task execution. We performed an open loop optogenetic manipulation, randomly delivering pulsed light (20 hz; 10 ms pulse width) for 1200 ms over 1/10^th^ of the behavior session (Figure 3K). When we examined wheel pulls similar to those executed during trials but where light was delivered before trial initiation (50-75ms before the peak of the pull), we observed a decrease in both velocity (p=0.1558 and p=0.0016 for control and hChR2, respectively, two-way ANOVA) and wheel pull distance (Figure 3L-N and Figure S8J; p=0.0708 and p=0.0011 for control and hChR2, respectively, two-way ANOVA). Together, these data reveal that activating M1^CT^ neurons at precise movement times when their activity normally decreases rapidly impairs movement execution regardless of the phase of movement, and strongly suggest that the decrease in activity of M1^CT^ neurons is permissive for this learned movement.

Our imaging experiments revealed that the activity of M1^CT^ neurons had lower magnitude anti-correlation to movement early in training relative to late training (Figure 2L, M). Consistently, our sc-RNAseq screen detected a gradual enrichment of this cell type with more enrichment in late training than in early training (Figure 1G). Therefore, we tested whether this activity signature was critical for forelimb movement early in training. As before, we set up a cohort of opsin and control FoxP2-cre animals but trained them to the early learning stage and delivered light concurrent with trial start, (20 hz; 10 ms pulse width; Figure 4A). Overall, the effect of this manipulation on movement execution was weaker with a non-significant decrease in the number of successful trials performed with light on in opsin expressing animals relative to control animals compared to the late training manipulations (Figure 4B,C; for percent of successful trials, p=0.0933, two-tailed unpaired t-test; for percent success, p=0.9771 and p=0.1021 for control and hChR2, respectively, two-way ANOVA). The most apparent deficit was observed when comparing how trials were executed (Figure 4D,E and Figure S9A,B,D,E) where we noted a decrease in wheel velocity in light trials only in hChR2 animals (for velocity trace: p<0.0001 and p>0.9999 for opsin and control, respectively; two-way ANOVA for 400ms after light on, time x condition; for mean max velocity: p=0.9810 and p=0.0331 for control and hChR2, respectively, two-way ANOVA) although this did not extend to total pull distance (Figure 4F and Figure S9C,F; p=0.5918 and p=0.1909 for control and hChR2, respectively, two-way ANOVA).

**Figure 4.**
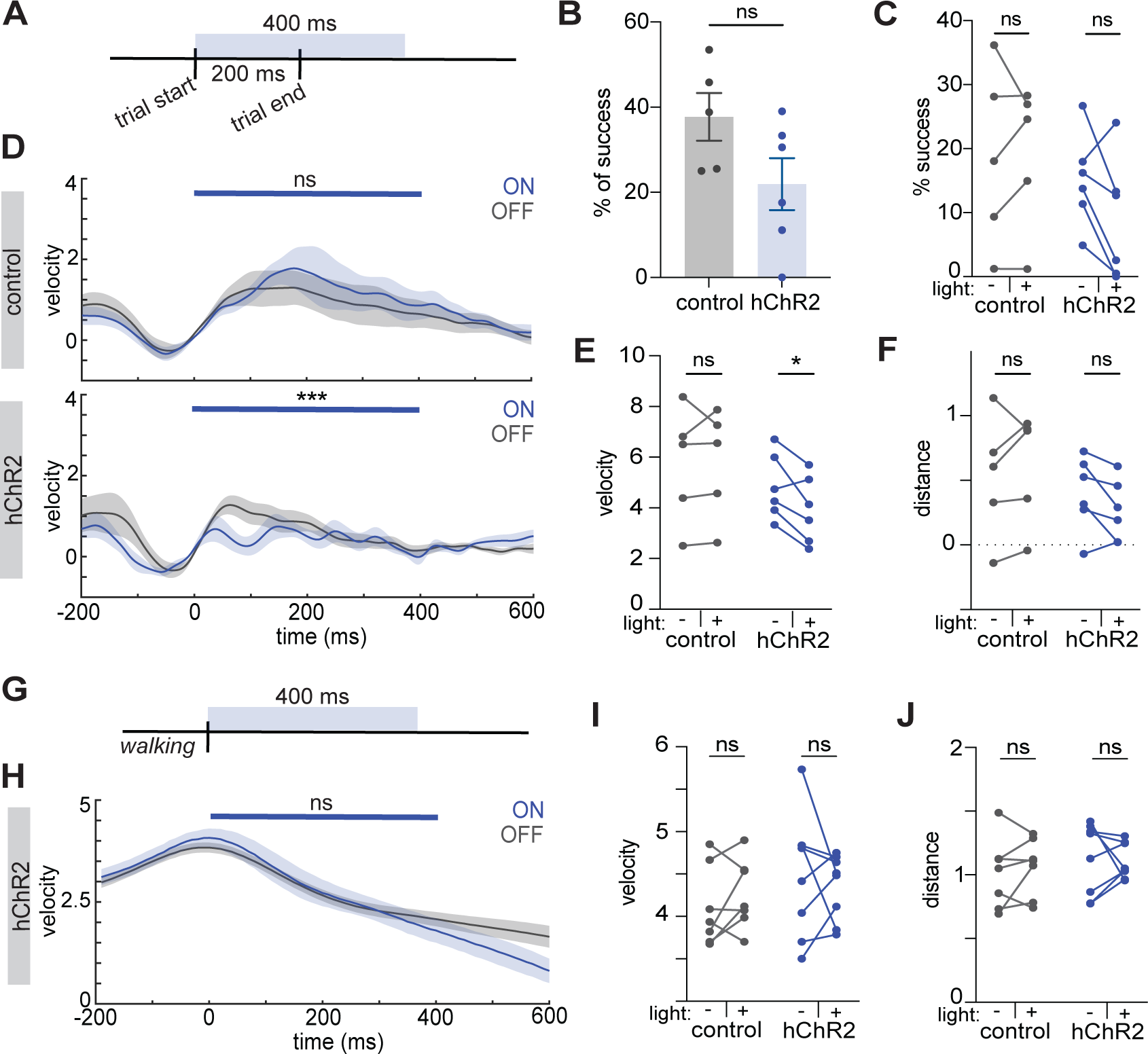
Differential perturbation to movement during manipulations of M1^CT^ activity. **A** Schematic of optogenetic light delivery for closed loop at trial start for wheel turning task at early training stage. Light was pulsed at 20 hz; 10 ms pulse width. **B** Percent of successful trials with light on (# successful trials during light on / # of all light on trials) control, gray; hChR2, blue; error bars, SEM; two-tailed unpaired t-test, p=0.0933. For B-J: N=5, control group; N=6, hChR2 group. Each point is average of all light delivery sessions. **C** Percentage of trials that are successful for each animal with light on or off in each group, (# (un)successful trials during light on / # of all (un)successful trials); two-way ANOVA, multiple comparisons; p=0.9771, control; p=0.1021, hChR2**. D** Velocity traces of all trials from control, top, and hChR2, bottom, animals during light, blue, and no light trials, gray; 0=trial start; blue bar denotes time of light, p>0.9999 (control), p<0.0001 (hChR2), time x condition, two way ANOVA; solid line, mean; shaded area, SEM. **E** Maximum velocity of all trials for each animal with light on or off in each group; two-way ANOVA, multiple comparisons, p=0.981 (control), p=0.0331 (hChR2). **F** Wheel distance traveled during all trials for each animal with light on or off for each group; two-way ANOVA, multiple comparisons; p=0.5918, (control), p=0.1909 (hChR2)**. G** Schematic of closed-loop optogenetic light delivery trials during walking bouts. **H** Velocity traces of walking trials from hChR2 animals (see Figure S10A for control) during light, blue, and no light trials, gray; 0=trial start; blue bar denotes time of light, p>0.9999 (control), p=0.9958 (hChR2), time x condition, two way ANOVA; solid line, mean; shaded area, SEM. **I** Max velocity during walking trials for each animal with light on or off in each group; two-way ANOVA, multiple comparisons, p=0.6981(control), p=0.7647 (hChR2). **J** Distance traveled during walking trials for each animal with light on or off in each group; two-way ANOVA, multiple comparisons, p=0.7998 (control), p=0.9605 (hChR2).

### Significance of M1^CT^ suppression is dependent on type of movement

We next investigated if the activity of M1^CT^ neurons had the same suppression during a more naturalistic movement that does not require extensive training. We decided to focus on locomotion because a) it is a movement that engages the forelimb in a different biomechanical sequence as well as many other muscle groups to generate a whole-body movement, and b) circuits in M1 as well as downstream circuits in the midbrain and hindbrain that mediate locomotion are distinct from those that are engaged in task specific forelimb movement^12,50^ providing an interesting comparison for our forelimb reaching task. We utilized the same two-photon imaging approach as before to record calcium dynamics in the dendrites of FoxP2+ M1^CT^ neurons. The predominant population still consisted of downmodulated neurons as in the wheel pulling task although at a somewhat lower proportion (Figure S10B,C). To examine if the suppression of M1^CT^ activity is critical for walking, we performed similar closed-loop optogenetic perturbations of FoxP2-cre animals expressing either a control fluorescent protein or channelrhodopsin (hChR2) as head fixed mice performed self-initiated walking bouts on a running wheel. Light was delivered after mice had commenced a walking bout (20 hz; 10 ms pulse width; Figure 4G). Here, we observed no difference in how mice walked when comparing light on or off trials in either the opsin or control group (Figure 4H and Figure S10A; p=0.9958 and p>0.9999 for opsin and control, respectively; two-way ANOVA for 400ms after light on, time x condition), the max velocity reached during the light delivery period (Figure 4I, p=0.6981 and p=0.7647 for control and hChR2, respectively, two-way ANOVA), or distance traveled (Figure 4J, p=0.7998 and p=0.9605 for control and hChR2, respectively, two-way ANOVA).

### M1^CT^ regulation of M1 corticospinals scales with learning

Our findings that M1^CT^ neurons are suppressed during the execution of learned forelimb movements contrast with the positive correlation with movement reported for other output populations of motor cortex, notably corticospinal neurons of Layer V^27^. We confirmed the increased activity of M1 corticospinal neurons during our wheel turning task as well as locomotion by using a retrograde labeling approach from the cervical spinal cord (Figure S11A) and imaging the dendritic dynamics of this population through a cranial window. In both wheel turning and walking, the activity of this population was positively correlated with movement (Figure 5A,B), and the predominant population consisted of up modulated neurons (Figure S11C,D). Remarkably, we observed that the magnitude of this correlation with movement increased during training (Figure 5C, S11B), a pattern symmetric to what we observed for M1^CT^ neurons.

**Figure 5.**
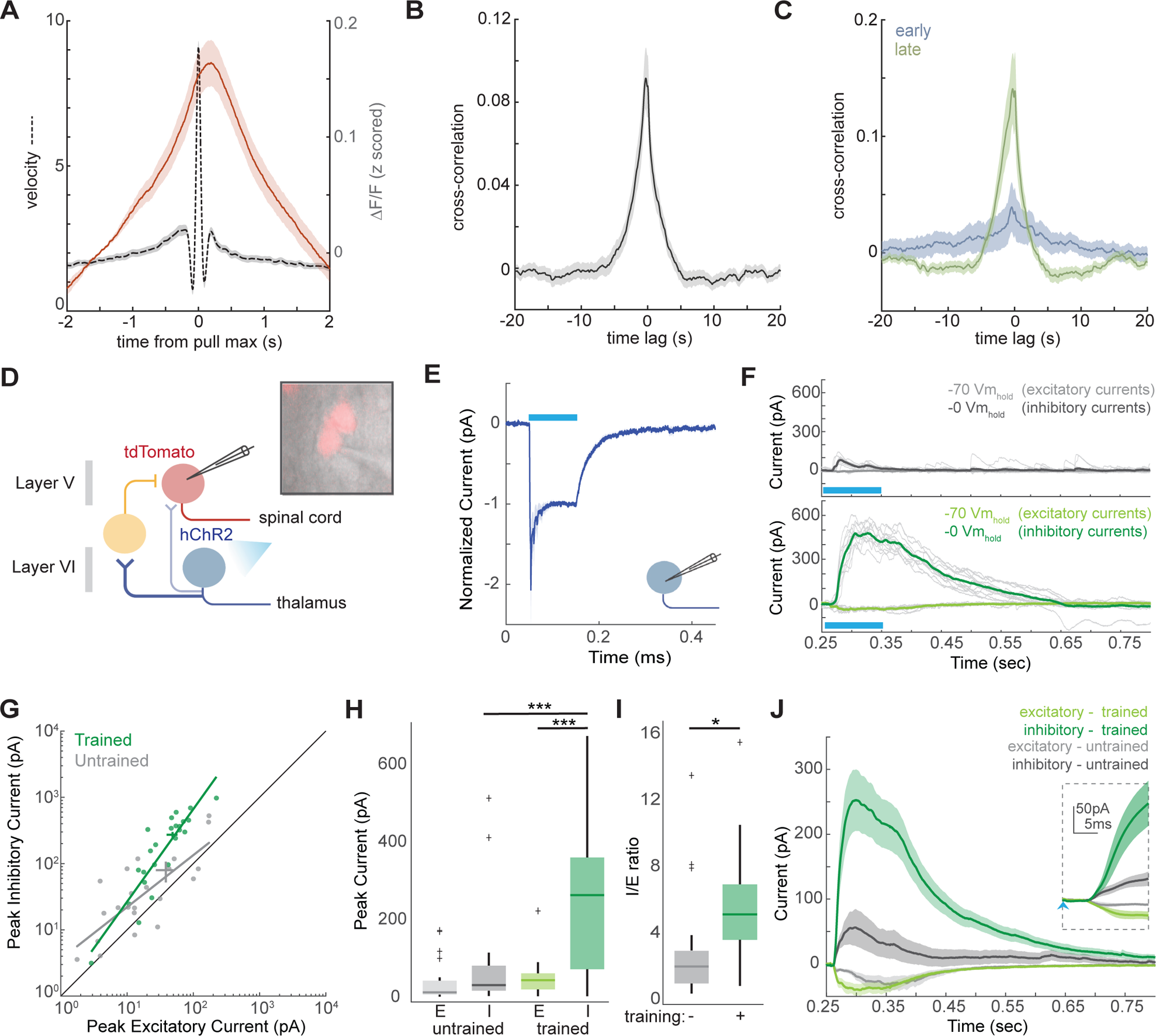
Multisynaptic inhibition of corticospinal neurons by M1^CT^ neurons augments with training. **A** Z-scored ΔF/F during wheel pulls; mean, solid red line; shaded area, SEM; Wheel velocity during same trials, dotted line; n=66 from N=3 animals; **B** Cross-correlation of Z-scored calcium ΔF/F to wheel velocity for whole session wheel pulls; mean, solid gray line; shaded area, SEM; **C** Cross-correlation to wheel velocity for pulls at early (blue) and late (green) training sessions; solid line, mean; shaded area, SEM. **D** Schematic of recording strategy from tdTomato labeled corticospinal neurons while activating M1^CT^ neurons expressing hChR; putative inhibitory neuron, yellow. **E** Population grand average response of corticothalamic neurons to photostimulation; light, blue bar; solid line, mean; shaded area, SEM, responses are normalized to the steady-state evoked current. **F** Recordings from exemplar corticospinal neurons after 100ms photostimulation (blue bar) from untrained, top, gray, or trained, bottom, green, animals at either −70mV holding potential, to measure excitatory currents, light gray and light green, or 0 mV holding potential, to measure inhibitory currents, dark gray or dark green; thick line, mean; thin lines, single trials. **G** Pairwise comparisons of peak excitatory and inhibitory currents from corticospinal neurons from untrained, gray, or trained, green, animals. Green and gray line, first polynomial linear fit, R^2^= 0.7303, untrained, R^2^= 0.7595, trained, simple linear regression **H,I** Comparison of peak excitatory (E) or inhibitory (I) currents **(H)** or inhibitory to excitatory ratio **(I)** from untrained, gray, or trained, green, animals. For **(H)** p<0.0001, peak inhibitory current in trained v untrained and p<0.0001 for trained peak inhibitory current v trained peak excitatory current, two-way ANOVA, multiple comparisons, see Table 4 for p values for all other comparisons; for **(I)**, p=0.0136, two-tailed unpaired t-test. **J** Grand average excitatory responses, light, or inhibitory responses, dark, from all neurons; mean, solid line; shaded area, SEM; inset for first 18 ms following the onset of photostimulation (indicated with blue arrow)

Given the opposing activity of these two populations, we hypothesized that M1^CT^ neurons could drive feedforward inhibition of corticospinal neurons in a similar circuit configuration to that reported in other cortical regions such as visual cortex^51,52^ and somatosensory cortex^53,54^. In this model, suppression of M1^CT^ neurons would release feedforward inhibition and allow for increased corticospinal activity. To address this possibility, we performed whole cell voltage-clamp recordings from fluorescent, retrogradely labeled corticospinal neurons in acute, live brain slices expressing Cre-dependent hChR2 in M1^CT^ neurons (Figure 5D and Figure S12A,B). Here, we utilized the Ntsr1-cre mouse to avoid any hChR2 expression in Layer V. We separately confirmed that photostimulation (100 ms) drove direct excitatory currents in M1 ^CT^ neurons (Figure 5E), as well as the baseline firing properties of M1^CT^ neurons (Figure S12D-F). We found hChR2 stimulation of M1^CT^ drove weak excitatory postsynaptic currents (EPSCs) in M1 corticospinal neurons when recording at membrane holding potential of −70 mV, but stronger inhibitory postsynaptic currents (IPSCs) at membrane holding potential of 0 mV (Figure 5F, top, G,H,J, gray). These findings are consistent with prior reports of disynaptic inhibition from M1^CT^ to corticospinal neurons^40^. Bath application of GABAzine eliminated IPSCs, confirming these inhibitory photocurrents are mediated by GABA_A_ receptors (Figure S12C, p=0.0284, paired t-test). Interestingly, we observed that the magnitude of this inhibition changed with training (Figure 5F,G,J) with a significant increase in the peak inhibitory current (Figure 5H, p<0.001, two-way ANOVA). This change corresponded with a higher inhibitory to excitatory ratio in corticospinal neurons with training, as the change in excitatory responses was less pronounced (Figure 5G, p=0.0136). However, we did note that excitatory and inhibitory responses became more coherent with training with higher consistency and shorter latency (Figure 5J, inset). Together, this led us to conclude that M1^CT^ neurons regulate the activity of corticospinal neurons through feedforward inhibition, whereby suppression of M1^CT^ neurons would result in disinhibition of corticospinal neurons. Finally, we observed that this feedforward inhibition was strongly facilitated with learning (Figure 5H).

## DISCUSSION

Our studies identified M1^CT^ neurons as a key permissive population for motor execution. We first identified this population using an activity-based, single-cell RNA-seq based screen of M1 during the progression of a motor task. Experiments characterizing the dynamics of M1^CT^ neurons showed that most of these neurons have a marked suppression during movement, and that this suppression is more substantial in late training than early training. Accordingly, closed-loop experiments revealed that optogenetic activation of these neurons during late training at different phases of movement execution rapidly and substantially impaired execution, suggesting that the decrease in activity is permissive for movement execution. However, closed-loop optogenetic activation of M1^CT^ neurons early in training or during locomotion had less impact on movement. Finally, we demonstrate that activity of M1^CT^ neurons suppresses the activity of M1 corticospinal neurons through feedforward inhibition, and that the strength of inhibition changes with learning. Our experiments uncover a mechanism by which suppression of activity of M1^CT^ neurons can disinhibit corticospinal neurons permitting them to fire during learned forelimb movements.

Our activity-based screen fills a niche in exploring cell type specific contributions to short time scale behaviors. Identifying the cell types with patterns of engagement during a particular behavior complements hypothesis-driven investigation of the role of particular cell types. This can then be followed by confirmatory techniques for neuronal activity recording, such as calcium imaging, which allow for in-depth characterization of particular activity patterns. Whereas we chose to focus on a cell type that differed in enrichment with training, it may also be informative to examine cell types that constitute the largest part of the active ensemble at a given point. In our experiments, we observed greater numbers of several IT subsets at the time points assessed. This could imply that these neurons are generally active for cortical processing regardless of the specific ongoing behavior. Although our screen has limitations and is not comprehensive, it still allowed us to compare differential cell type engagement during training. We also chose to deliver light unbiasedly through the training session to include all activity that could contribute to learning, not just movement execution, but groups interested in more specific features of a given behavior could refine periods of light delivery. As new tools emerge, they may allow for similar but more quantitative approaches for identifying active cell types in the future.

There has been a long-standing interest in defining the role of the primary motor cortex and its relationship to movement. Prior studies have highlighted the heterogenous patterns of activity relative to movement, but narrowing these examinations to specific cell types has begun to produce a more precise relationship between neural activity and movement. For example, we and others have noted that most M1 corticospinal neurons, which project directly to the spinal cord and are hence relatively proximal to skeletal muscle, have predominantly movement correlated activity^27^. Similarly, intratelencephalic neurons projecting within cortex show increased activity during movement^31^. In contrast, our imaging of M1^CT^ neurons shows that the majority of this population reduces activity at the time of movement. This signature has also been recently reported for corticopontine neurons of layer V^31^ highlighting the heterogeneity within M1 layers. While these examinations allow us to begin integrating the roles of these cell types during movement, it remains to be seen whether much more refined targeting of smaller subsets of neurons as defined by transcriptomic profiling will yield a more complete picture of the role of M1.

Our closed-loop optogenetic experiments suggest that activation of M1^CT^ neurons at a time when they should be suppressed can rapidly and profoundly impair movement execution. Our experiments further suggest that feedforward inhibition from M1^CT^ neurons corticospinal neurons, (Figure 5) which are positively correlated with movement and are thought to relay motor commands throughout the brain and spinal cord. The change we observed in the magnitude of inhibition with training is also reflected in the magnitude of movement perturbation observed in our optogenetic experiments at early or late training. This further highlighting this mechanism as a potential source of refinement that integrates the activity of these population during learning. It should also be noted that among the targets of corticospinals are many of the same thalamic regions as those targeted by layer VI corticothalamic neurons. We demonstrated that within layer V and VI, similar patterns of topographical organization occur withing M1 relative to thalamic targets (Figure S3) indicative of a columnar organization. This coupling of output targets underscores the relevance of local M1 mechanisms to regulate output signals from M1. In the future, it would be interesting to examine the activity of these thalamic targets regions and to investigate any additional mechanism for integrating these signals within both M1 and thalamus.

It is intriguing to note that a similar mechanism of local, disynaptic inhibition by layer VI neurons to all other layers has been described in visual and somatosensory cortices^51–54^. Our parallel observation in motor cortex brings up avenues of further exploration. For example, it would be informative to examine whether M1^CT^ neurons also drive a similar inhibitory input to more superficial layers within motor cortex, such as layer II/III which is thought to perform a more distinct role in processing inputs to M1 rather than relaying output signal as do deeper layers. Likewise, it will be interesting to explore more general functions of this modulation in M1. In visual cortex, the inhibitory modulation provided by layer VI neurons to superficial layers plays an important role in gain control. We observed a gradation in the suppression of M1^CT^ neurons relative to movement vigor (Figure 2H-K), suggesting a similar modulatory role for movement. If true, this could suggest a common role for this population and this circuit motif across different regions of cortex.

## ACKNOWLEDGEMENTS

We are grateful to C. L. Warriner for sharing a prior iteration of the wheel turning task; H. Rodrigues, for assistance designing and constructing behavioral equipment; D. Ng and D. Peterka for discussion and feedback on the manuscript; I. Shieren, G. Martins, and M. Correia for technical assistance; L. Hammond, D. Peterka, and H. Ibarra Avila for assistance with various types of imaging and imaging data analysis; and T.M. Jessell for invaluable enthusiasm and support when this project was devised. Imaging was performed with support from the Zuckerman Institute’s Cellular Imaging platform, and the National Institute of Health (NIH 1S10OD023587-01). L.M.C. was a Hellen Hay Whitney Foundation Fellow and is currently supported by a NIH Pathway to Independence Award (1K99NS127857-01). R.M.C. was funded by grants from the NIH (5U19NS104649) and the Simons-Emory International Consortium on Motor Control.

## AUTHOR CONTRIBUTIONS

L.M.C., A.N., R.M.C., and V.M. designed experiments and interpreted data; L.M.C. and A.N. performed experiments and analyzed data; L.T. collected and analyzed anatomical data; A.K. assisted in experimental optimization; R.S. assisted with tissue clearing; and M.D.K performed all flow cytometry; and L.M.C and R.M.C. wrote the manuscript.

## MATERIALS AND METHODS

### Mice

All experiments and procedures were performed according to National Institutes of Health (NIH) guidelines and approved by the Institutional Animal Care and Use Committee of Columbia University. Adult mice of both sexes, aged 2–6 months, were used for all experiments, except for RNA sequencing which only used males aged 3 months. The strains used were: C57BL6/J (Jackson Laboratories, 000664), FoxP2-Cre (Jackson Laboratories, 030541), and Ntsr1-Cre (Mutant Mouse Resource & Research Centers, 030648-UCD). Only mice used for RNA sequencing experiments were individually housed. All mice were kept under a 12-h light-dark cycle. All mice except those used for calcium imaging experiments were kept under reverse light cycle conditions.

### Viral Vectors

For CaMPARI based experiments, AAV1-hSyn-CaMPARI (Addgene 100832-AAV1) was used. Projection mapping was performed with the membrane bound GFP expressed from AAV8-hSyn-JAWS-KGC-GFP-ER2 (Addgene 65014-AAV8). Calcium imaging experiments were performed with AAV1-Syn-FLEX-jGCaMP7f (Addgene 104492-AAV1) for M1^CT^ neurons or AAVretro-hSyn-FLEX-jGCaMP7f (Addgene 104492-AAVrg) in combination with AAV1-hSyn-Cre (Addgene 105553-AAV1) for corticospinal neurons. Optogenetic experiments used AAV1-Ef1a-DIO-EYFP (Addgene 27056-AAV1) and AAV1-EF1a-DIO-hChR2 (H134R)-EYFP (Addgene 20298-AAV1) for control and opsin animals respectively. Slice electrophysiology experiments used AAV1-EF1a-DIO-hChR2 (H134R)-EYFP (Addgene 20298-AAV1) injected in M1 and AAVretro-CAG-TdTomato (Addgene 59462-AAVrg) injected into spinal cord for photostimulation experiments. Baseline recordings used AAV5-hSyn-DIO-mCherry (Addgene 50459-AA5).

### Stereotactic surgery

Animals were anesthetized using isofluorane, and analgesia was delivered subcutaneously in the form of either carprofen (5 mg/kg) or burprenorphine SR (0.5-1 mg/kg) as well as bupivacaine (2 mg/kg). For all viral vectors delivered to M1, injections were centered in the caudal forelimb area (CFA) using the following coordinates: 1.5 mm lateral to the midline, 0.25 mm rostral to bregma. A total of 5 injections were made in this region to cover a large portion of CFA in the contralateral (left) hemisphere. One injection was at the center of these coordinates, with 4 additional injections either 300 um anterior/posterior to this or 300 um medial/later to this. Specific sites were adjusted to avoid blood vessels. A Nanoject III Programmable Nanoliter Injector (Drummond Scientific) was used at a rate of 1 nl/s.

For CaMPARI experiments, a total of 1 uL of virus was delivered (200 uL at each of the five injection sites) at a range extending to 800 um below the pial surface (25 uL every 100 um). For projection mapping, a total of 500 uL of viral vector was injected (100 nL per injection site) from 900-500um below the pial surface (20 nL at every 100 um). For calcium imaging of M1^CT^ neurons, a total of 375 uL of viral vector was injected (75 nL per injection site) from 950-650 um below the pial surface (25 nL at every 100 um). In FoxP2-cre mice or for imaging corticospinal neurons, a total of 1uL was injected into the right cervical spinal cord (200 nL per segment from C3-C7). For optogenetic experiments, a total of 500 uL of viral vector was injected (100 nL per injection site) from 850-650um below the pial surface (20 nL at every 100 um). For photostimulation slice recordings, a total of 240 nL was delivered (60 nL at each of 4 injection sites) at a range extending from 650-850 um below the pial surface (20 nL every 100 um) along an injection in the right cervical spinal cord of a total of 1 uL (200 nL per segment from C3-C7). For baseline recordings, a single injection of 50 nL of diluted virus (2.1×10^12^ GC/mL) was delivered at 800 um below the pial surface. To prevent backflow, we waited 3-5 minutes before the initial injection and 5-10 minutes after each injection. For animals used in behavioral experiments, a headplate was attached to the skull with Metabond (Parkell) post injection.

For retrograde labeling from thalamus, we used the following coordinates, for the ventral region: 1.3 mm lateral from the midline, 1.22 mm caudal to bregma, and 3.5 mm below the pial surface; for the lateral region: 1.25 mm lateral from the midline, 2.18 mm caudal to bregma, and 3.25 mm below the pial surface; for the medial region: 0.1 mm lateral from the midline, 1.46 mm caudal to bregma, and 4.2 mm below the pial surface. A total of 10 nL of 4% FluoroGold (Fluorochrome) was injected. FluoroGold was freshly diluted the day of injection from a 10% stock solution. To prevent backflow, we waited 2 minutes before each injection and 10 minutes after each injection.

### Cranial window implantation

For animals requiring a cranial window, surgery was performed as described above with the following modification. Animals were anesthetized using isofluorane, and analgesia was delivered subcutaneously in the form of burprenorphine SR (0.5-1 mg/kg) and bupivacaine (2 mg/kg). Dexamethasone (2 mg/kg) was also delivered as an anti-inflammatory agent. A larger craniotomy was made (2.5 mm in diameter) for a custom cranial window made of a glass plug (2.5 mm diameter) attached to a larger glass base (3.5 mm in diameter) with optical cement (Norland Optical Adhesive 61). After viral injection, the cranial window was implanted to gently press on the brain at the site of the craniotomy and secured with Metabond (Parkell). Headplate implantation followed cranial window implantations.

### Behavior

For the wheel pulling task, several noise attenuating behavioral chambers equipped with IR light were assembled with custom components to train several animals in parallel. Mice were placed in flat bottom, opaque tube, and head fixed using a custom metal headplate by screwing in place to adjustable head posts. The small rotary wheel was directly below the right forelimb of the animal when in the holder. A small partition prevented animals from utilizing their left forelimb to pull the wheel. A small screw was also provided for left forelimb placement. Mice were habituated to head fixation and water delivery via the water port for several days before the beginning of training. The water port consisted of a blunt needle positioned in front of the mouse so water was reachable by licking. Water droplets were generated using a solenoid valve and were calibrated before the beginning of training for each experiment. The 60 mm diameter rotary wheel was assembled from two acrylic sides held together by small metal screw rungs. An absolute rotary encoder (US Digital; 10 bit) was placed in the shaft of the wheel to assess wheel movement. Each rung was wired to a capacitance sensor. An IR beam was placed at the apex of the wheel on a custom 3D printed holder allowing for alignment of the beam. Data from all behavioral components was collected using a DAQ (National Instruments) sampling at 2000 Hz. Behavioral sessions were controlled using custom MATLAB code which assessed beam breaks and changes in rung capacitance. If thresholds were met, a trial would commence and wheel displacement was then assessed, smoothed, and converted to velocity. If the threshold velocity was surpassed, a pulse was sent to the solenoid to deliver the water reward. Mice were habituated to the task during three pretraining sessions prior to the beginning of training. During these sessions, mice were exposed to the task but at low trial velocity thresholds for a limited number of trials. Training sessions consisted of 30 minutes daily or a maximal of 100 successful trials. Mice were trained during the dark phase of their reversed light cycle housing schedule.

For locomotion, mice were headfixed atop a large acrylic rotary wheel places inside a noise attenuating chamber. The wheel was lined with clingwrap for traction. Mice were habituated to head fixation before the beginning of training. A quadrature rotary encoder (US Digital; 1024 CPR) was placed in the shaft of the wheel to assess wheel movement. The behavioral assays were controlled and data was collected using pyControl.

### CaMPARI labeling

Four weeks post injection, mice were head-fixed as in all behavioral experiments and a plastic fiberoptic patch cord (960 um core; 0.63 NA) ending in a metal ferrule (Doric) was placed directly over the cranial window. A multichannel LED driver (Doric) was used with 405nm central wavelength connectorized LED (Doric, ∼60 mW). Concurrent with the start of the training session, light delivery was initiated using the Doric Neuroscience Studio software. Light was delivered in square wave pulses of 5s each with 5s of light off between each pulse for a total of 12 minutes.

### Generation of single-cell suspensions and FACS

After CaMPARI labeling, mice were returned to their home cage for 15 minutes and then euthanatized by transcardial perfusion with ice-cold choline solution as previously described^55^. Thick sections were cut using a matrix, and M1 was quickly dissected. Single-cell suspensions were generated using the previously described protocol optimized for preserving transcriptional state with the following modifications^55^. The dissected M1 tissue was further triturated with spring scissors. Incubation with papain was conducted using a dialysis membrane (Fisher Scientific, Slide-a-Lyzer MINI Dialysis Device,10k molecular weight cut-off) suspended in a beaker of dissociation solution allowing for constant oxygenation (95% O2 and 5% CO2) of the solution without disrupting enzyme integrity within the dialysis membrane. After enzymatic dissociation, the tissue was broken into a single-cell suspension by gently pipetting using wide-bore pipette tips. The sample was cleaned using the Debris Removal Solution (Miltenyi Biotec) as described by the manufacturer. Cells were suspended in dissociation solution containing 0.04% BSA, Vybrant DyeCycle Ruby for exclusion debris, and RNAase inhibitors. Cell sorting was performed using a FACSAria Cell Sorter (Becton Dickinson) with a 130 um nozzle at 12 PSI using the following laser lines and filters: CaMPARI Green, 488 nm, 530/30 bandpass; CaMPARI Red, 561 nm, 610/20 bandpass; DyeCycle Ruby, 637 nm, 670/30 bandpass. Gates were set using a sample not expressing CaMPARI and a sample not exposed to photoconversion light. Any cells with detectable red fluorescence were collected.

### Single-cell RNA-seq

Sorted cells were captured and barcoded using 10x Genomics Chromium v3 according to the manufacturer’s protocol. Samples were processed and libraries were prepared and sequenced by the JP Sulzberger Columbia Genome Center Single Cell Analysis Core.

### Single-cell RNA-seq analysis

Single-cell RNA-seq analysis followed a standard workflow, comprising the following steps:

1. Sequencing read files were aligned to the mm10 genome using 10x Genomics Cellranger v.3.1.0, generating a table of cell barcodes by Unique Molecular Identifier (UMI) counts per gene.
2. Raw counts files were aggregated, and cells with >20% UMIs mapping to mitochondrial genes were removed, as well as cells with <200 UMIs after removing mitochondrial genes, ribosomal protein genes, pseudogenes, and gene models.
3. Cells were clustered using the Seurat R package (v.4.0.0), with SCTransform (default parameters) and PCA, followed by integration across batches using the Harmony R package (with 40 PCs). The cell-cell neighbor network was constructed with 30 Harmony dimensions, and the Louvain community detection algorithm was run with resolution = 0.4.
4. Based on expression of marker genes for neurons and glia (Snap25, Slc17a6, Slc17a7, Gad1, Gad2, Mbp, Mog, Fgfr3, Aqp4, Pdgfra, Tmem119, Aif1, Ptprc, and Cldn5), the clusters corresponding to glutamatergic and GABAergic neurons were extracted.
5. Step 3 was re-run on the glutamatergic and GABAergic neurons (separately), with 30 PCs, 20 Harmony dimensions, and resolution = 0.2.
6. Combinatorial marker genes were identified for each cluster using the FindMarkers command in Seurat, run on all pairs of clusters. In parallel, putative layer identities for each cluster were assigned by mapping to Azimuth Mouse Motor Cortex reference.
7. To assess differential proportions among conditions, cell count values were run through the ANCOM-BC and MASC R packages, with pairwise comparison between control, early, and late conditions. All q-values were then further corrected.

### Projection mapping

Mice were euthanized by intracardial perfusion with 1× PBS followed by 4% paraformaldehyde. Brains were post-fixed in 4% paraformaldehyde overnight at 4 °C. The AdipoClear protocol was performed as previously described^56^ with the following modifications. Primary and secondary antibody incubations were conducted at 37 °C and for 6 days per incubation. The following primary antibodies were used at a dilution of 1:2500: chicken GFP antibody (Aves) and rabbit RFP antibody (Rockland). The following secondary antibodies were used at a dilution of 1:2500: donkey anti-rabbit Alexa Fluor 647 (ThermoFisher Scientific) and donkey anti-chicken Alexa Fluor 647 (ThermoFisher Scientific). Samples were imaged with a light-sheet microscope (Ultramicroscope II, LaVision Biotec) using a 4x objective for the fluorescent proteins or 1.3x objective for autofluorescence imaging. The ClearMap pipeline was used as previously described to map the data to the Allen Brain Reference Atlas^57^. All signal mapped to thalamus was summed and the percent of signal in each nucleus was calculated for each animal.

### Retrograde mapping

Two weeks post injection, mice were euthanized by intracardial perfusion with 1× PBS followed by 4% paraformaldehyde. Brains were post-fixed in 4% paraformaldehyde overnight at 4 °C. Serial coronal sections (50um) were collected and stained with NeuroTrace 640/660 (ThermoFisher Scientific). Imaging was performed with an AZ100 automated slide scanning microscope using a 4x objective (Nikon, 0.4 NA). Image processing, registration to the Allen Brain Reference Atlas, and automated counting were performed as previously described^58^. Cell counts per animal were summed over anatomical regions or binned according to their mapped position for comparative analysis.

### Two-photon imaging and analysis

All behavioral training and habituation were conducted as described above. Training did not commence until at least 8 weeks post viral injection. A behavioral assembly similar to those described for behavior was placed on a modified 2p microscope (Bruker) for calcium imaging experiments. A 25x water immersion objective (Olympus,1 NA) was used with a mode-locked Ti:sapphire laser (Verdi 18W, Coherent) at 920 nm. Images were collected with Prairie View software (Bruker) at 64 Hz and averaged every 4 images for an effective sampling rate of 16 Hz. The pial surface of the brain surface was identified at the beginning of each imaging session, and fields of view were selected at least 300um below the pial surface. Voltage recordings of the encoder and solenoid were also collected with Prairie View. The behavioral code was run from a separate computer, and all behavioral components recorded. The encoder signal was used to align the imaging to the behavioral recordings. Motion correction and signal extraction was performed using CNMF as previously described^48^ with an autoregressive process *p* of 2. For analysis of the whole session, the Δ*F/F* or deconvolved events were z-scored over the whole session for individual neurons and then aggregated across animals for further analysis. For analysis of trials or pulls, the Δ*F/F* or deconvolved events were z-scored to signal 2-1.5 s prior to the movement event for each neuron. ROC classification was performed by calculating the approximate integral using the trapezoidal method for 100 ms bins starting 2s before pull max and extending 2s after and z-scored to the first five bins (−2 to −1.5 from pull max). Neurons with at least two bins of the 3 proceeding and the 3 preceding the pull max (total of 600 ms) with a z-score greater than 2 or −2 were classified as up or down respectively. All other neurons were classified as not modulated.

### Optogenetic manipulations

All behavioral training and habituation were conducted as described above. On the days of optogenetic manipulations, a plastic fiberoptic patch cord (960 um core; 0.63 NA) ending in a metal ferrule (Doric) was placed directly over the cranial window. A multichannel LED driver (Doric) was used with 465nm central wavelength connectorized LED (Doric; ∼35 mW). For closed-loop experiments, a pulse was sent at trial start on one third of trials to the driver to trigger light delivery as programmed in Doric Neuroscience Studio software. Light was delivered for 400 ms at 20hz with a pulse width of 10ms. For open-loop experiments, light delivery was triggered using a random number generator to give light for 10% of the session. Light was delivered for 1.2 s at 20 hz.

### Slice Electrophysiology

Mice were deeply anesthetized with isoflurane and transcardially perfused with an ice-cold carbogenated solution of HEPES-sucrose artificial cerebrospinal fluid (ACSF) containing 110 mM NaCl, 10 mM HEPES, 25 mM glucose, 75 mM sucrose, 7.5 mM MgCl_2_, and 2.5 mM KCl. The brain was removed from the skull and glued to the stage of a vibrating microtome (Leica). Next, 300-μm coronal brain slices were cut in a bath of ice-cold, slushy, HEPES-sucrose ACSF. Slices were incubated for 30 min in a 34 °C bath of normal carbogenated ACSF containing 124 mM NaCl, 2.7 mM KCl, 2 mM CaCl_2_, 1.3 mM MgSO_4_, 26 mM NaHCO_3_, 1.25 mM NaH_2_PO_4_, 18 mM glucose and 0.79 mM sodium ascorbate. Slices were then transitioned to room temperature, where they remained for the duration of the experiment. Patch electrodes (1–3 MΩ) were filled with a cesium/QX-314-based internal solution containing 5 mM QX-314, 2 mM ATP magnesium salt, 0.3 mM GTP sodium salt, 10 mM phosphocreatine, 0.2 mM EGTA, 2 mM MgCl_2_, 5 mM NaCl, 10 mM HEPES, 120 mM cesium methanesulfonate and 0.15% Neurobiotin. All recordings were made using a Multiclamp 700B amplifier, the output of which was digitized at 10 kHz (Digidata 1440A). Series resistance was always <20 MΩ and was compensated up to 90%. Neurons were targeted with DIC microcroscopy and epifluorescence when appropriate. Excitatory or inhibitory currents were isolated by clamping membrane voltage at −70mV or 0mV, respectively. GABA_A_ currents were blocked by superfusion of 10uM Gabazine for 10 minutes. Neurobiotin-filled cells were visualized post hoc through streptavidin processing. Brain slices were fixed in 4% paraformaldehyde for 1 hour and rinsed several times in PBS. Slices were initially permeabilized in 1% Triton-containing PBS for 1 hour at room temperature. Slices were incubated overnight at 4C in 0.3% Triton PBS containing 0.1% Streptavidin DyLight Fluor 405, followed by several PBS rinses at room temperature.

### Histology and imaging

Images of CaMPARI photoconversion were obtained from mice trained as described above. Images of GCaMP expression were obtained from mice used for imaging once all training was complete. All mice were euthanized by intracardial perfusion with 1× PBS followed by 4% paraformaldehyde. Brains were post-fixed in 4% paraformaldehyde overnight at 4 °C. Coronal sections (50um) were cut with a vibratome. DAPI was used as a counterstain. GCaMP signal and hChR2-EYFP expression was amplified with chicken GFP antibody (Aves). TdTomato expression was amplified with rabbit RFP antibody (Rockland). For CaMPARI slices, imaging was performed with a confocal microscope (Zeiss 880) with a 20x objective within a week of mounting. All other images were acquired with a W1-Yokogawa spinning disk confocal microscope with a 4x objective for zoomed out panels or 20x objective for zoomed in panels.

### Statistical Analysis

Statistical parameters, statistical tests, and statistical significance are reported throughout. Significance is defined as P<0.05 with significance annotations of *P < 0.05, **P < 0.01, ***P < 0.001 and ****P < 0.0001. All statistical analysis was performed using R, GraphPad Prism, Python, or MATLAB.

### Data Availability

Single-cell RNA-sequencing data will be deposited in the Gene Expression Omnibus. All other data that support the findings of this study are available from the corresponding authors upon reasonable request.

### Code Availability

Custom code used in this study is available from the corresponding authors upon reasonable request.

---

**Figure S1.**
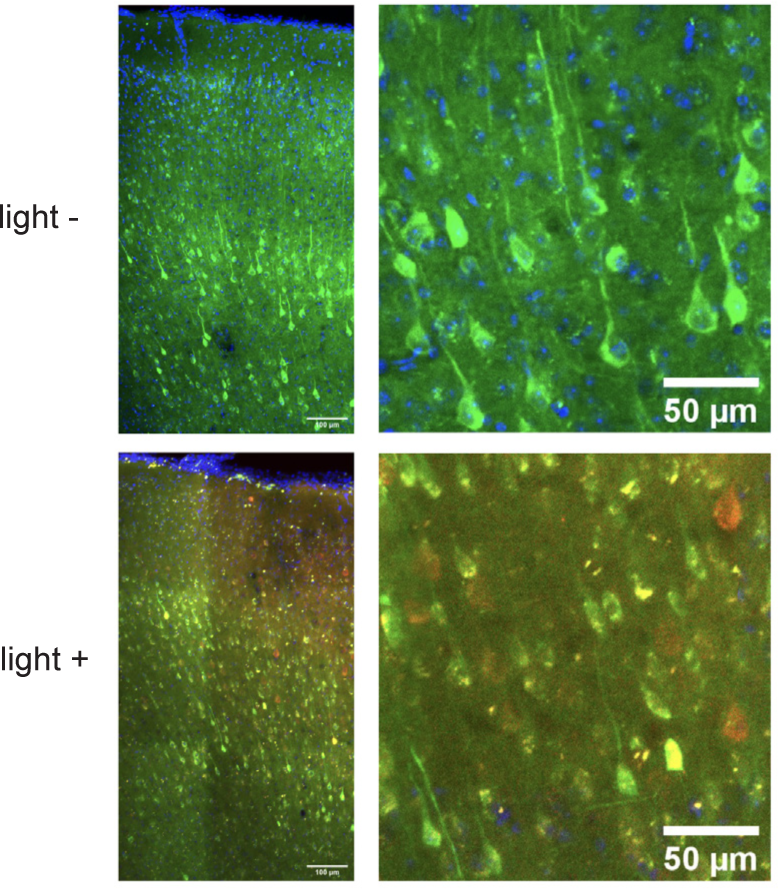
CaMPARI expression in M1 neurons. Representative images of CaMPARI levels in mice performing the wheel turning task in the absence (top) and presence (bottom) of photoconversion light. Green, baseline; red, photoconverted state. Right panels are zoom insets of left.

**Figure S2.**
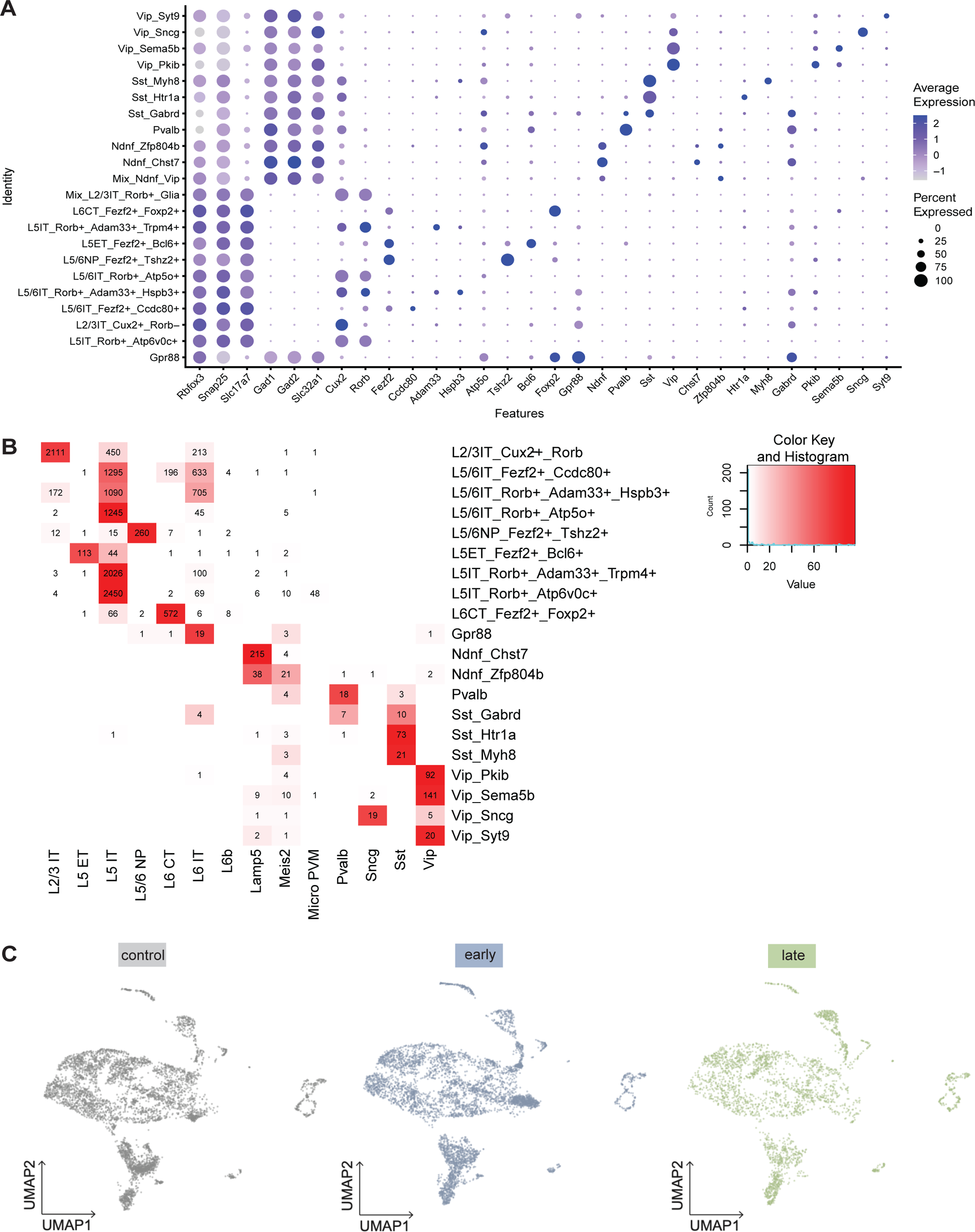
Gene expression and mapping of single cell RNA sequencing. **A** Average, purple gradient, and percent expression, size of dot, of marker genes in each cluster. **B** Number of cells mapped from clusters identified in aggregate sc-RNAseq runs to clusters of the M1 cell type atlas. **C** UMAPs of scRNA-seq runs from each condition mapped to M1 cell type atlas; gray, control; blue, early training; green, late training.

**Figure S3.**
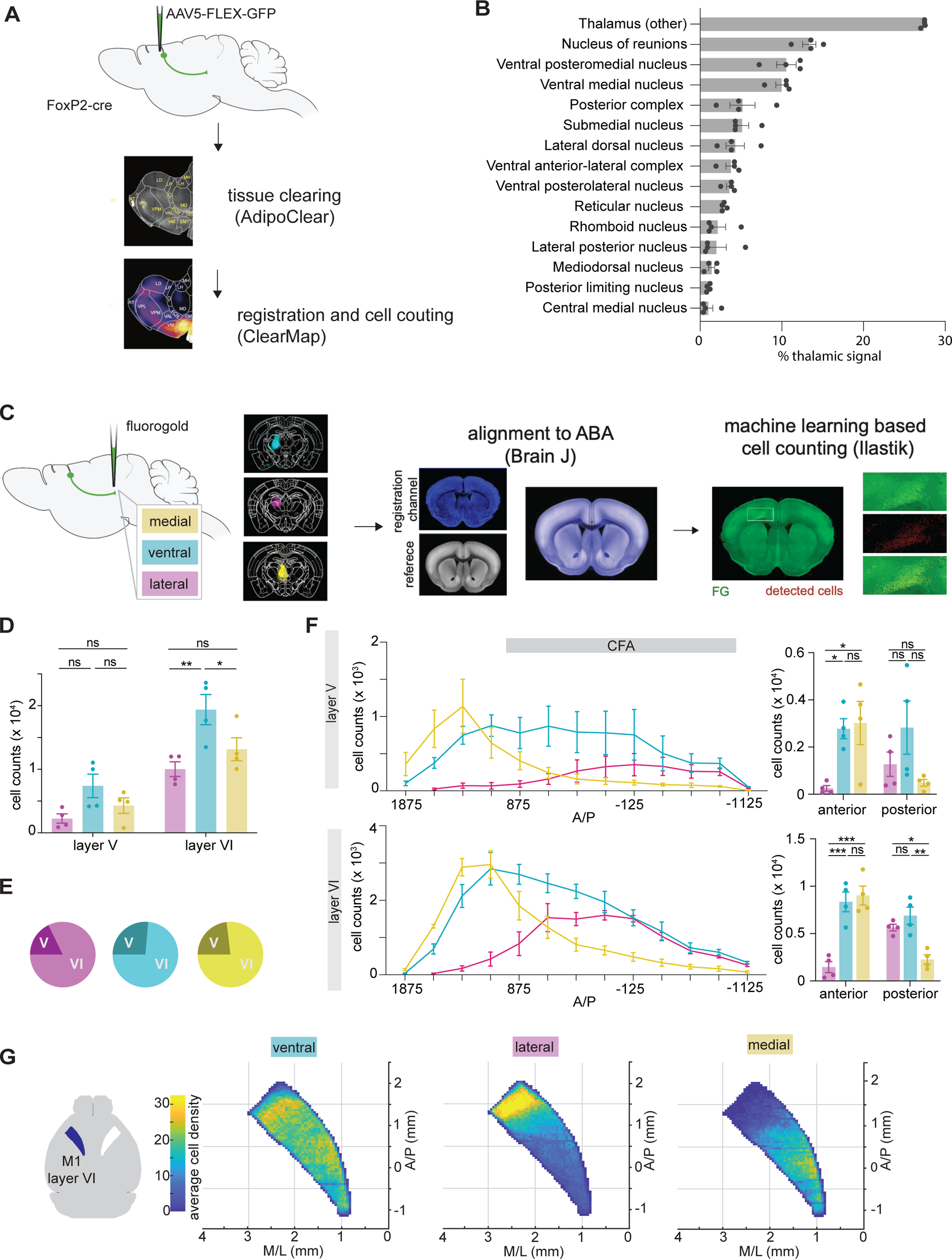
Topographical organization and projection pattern of M1^CT^ neurons. **A** Schematic of M1^CT^ projection mapping approach in thalamus from M1 in FoxP2-cre mice. Max projection of representative M1^CT^ projections in thalamus; mCherry labeling. Heatmap of thalamic signal; N=4. **B** Thalamic regions with greater than 1% of total thalamic signal; N=4; error bars, SEM. **C** Schematic of retrograde labeling of M1^CT^ neurons from three regions of thalamus: medial (yellow), ventral (cyan), and lateral (magenta). Heatmap of injection site, N=4 animals for all analysis (C-G) and color corresponds to region targeted (C-G). Subsequent workflow consisted of alignment and segmentation using BrainJ and Ilastik. **D** Number of M1^CT^ neurons labeled from each thalamic region in both M1 layer V and layer VI, two-way ANOVA, p=0.0020 (lateral v. ventral, layer VI); p=0.0400 (ventral v. medial, layer VI). **E** Percent of total signal in either cortical layer; **F** Left, Number of M1^CT^ neurons labeled in layer V (top) or layer VI (bottom) from each thalamic region along the anterior/posterior axis; 0=Bregma; error bars, SEM. Right, aggregate cell counts of M1^CT^ neurons labeled from each thalamic target in anterior (1625 to 875 um A/P) or posterior (125 to −375 um A/P) regions of M1; error bars, SEM; for layer VI: two-way ANOVA, p<0.001 (anterior, medial v. lateral); p=0.0173 (posterior, medial v. lateral); see Table 5 for all comparisons. **G** Distribution of cell density of layer VI M1^CT^ from each thalamic target; left, schematic of M1 layer VI, left hemisphere, blue, mapped.

**Figure S4.**
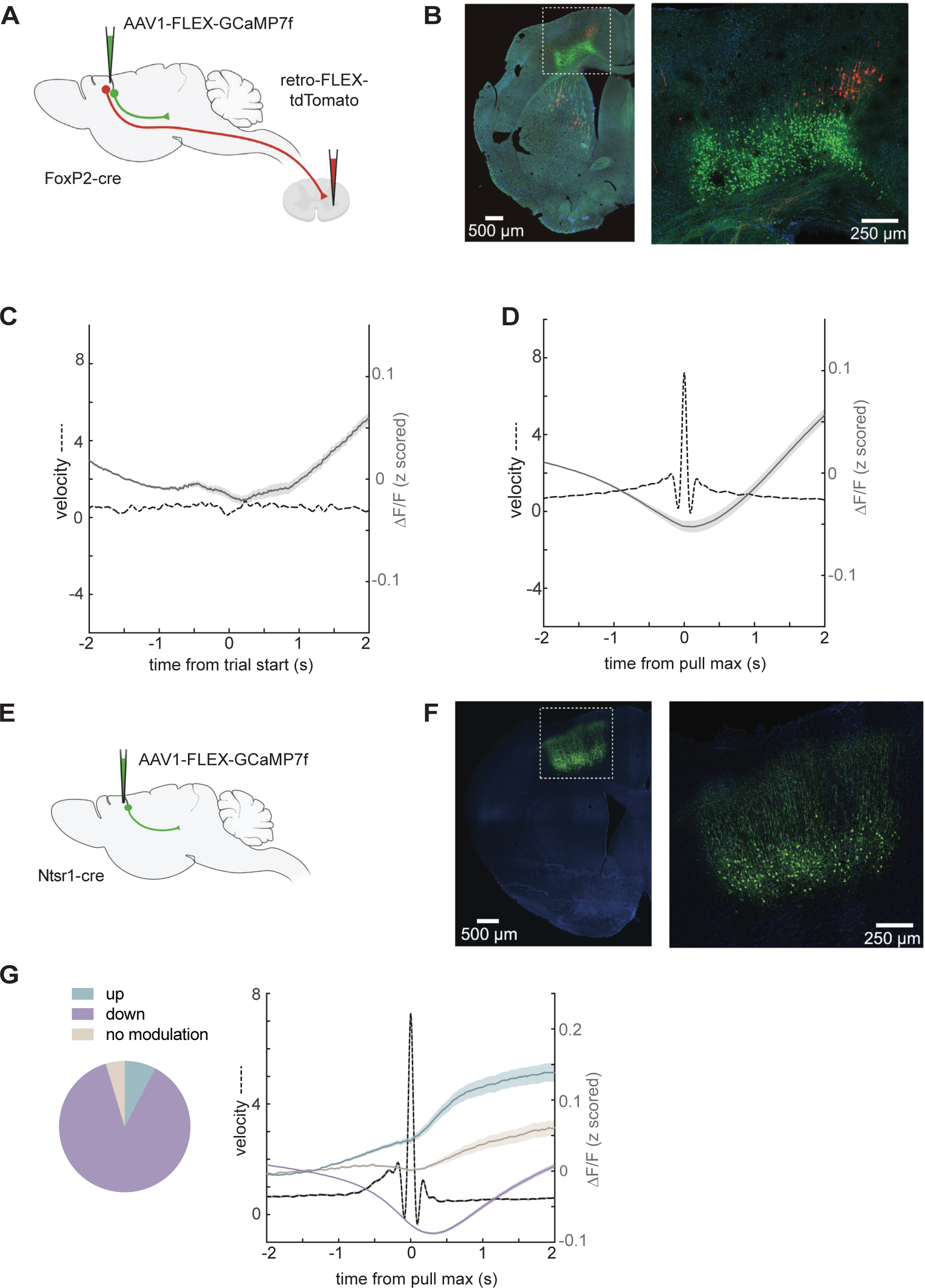
Calcium imaging approach and M1^CT^ activity. **A** Schematic of injection strategy for calcium imaging of M1^CT^ neurons in FoxP2-cre mice. An AAV expressing cre-dependent GCaMP7f was injected into the caudal forelimb area of M1. A retrograde AAV expressing cre-dependent tdTomato was also injected into the cervical spinal cord to allow for exclusion of FoxP2 expressing corticospinals from imaging fields. **B** antibody amplified GCaMP7f signal, green; DAPI, blue; corticospinal neurons, red. **C,D** Z-scored ΔF/F during unsuccessful trials **(C)** OR wheel pulls **(D)** mean, solid gray line; shaded area, SEM; Wheel velocity, dotted line; n=1078; N=4. **E** Schematic of injection strategy for calcium imaging of M1^CT^ neurons in Ntsr1-cre mice. **F** antibody amplified GCaMP7f signal, green; DAPI, blue; corticospinal neurons, red. **G.** Left, ROC based classification of all units across sessions based on activity during wheel pulls in Ntsr1-cre animals. Right, Z-scored ΔF/F of each ROC group during wheel pulls; solid lines, mean; shaded areas, SEM; wheel velocity during pulls, dotted line; 0=pull velocity max; n= 1052 downmodulate units, n=93 upmodulated units, n=56 no modulation units.

**Figure S5.**
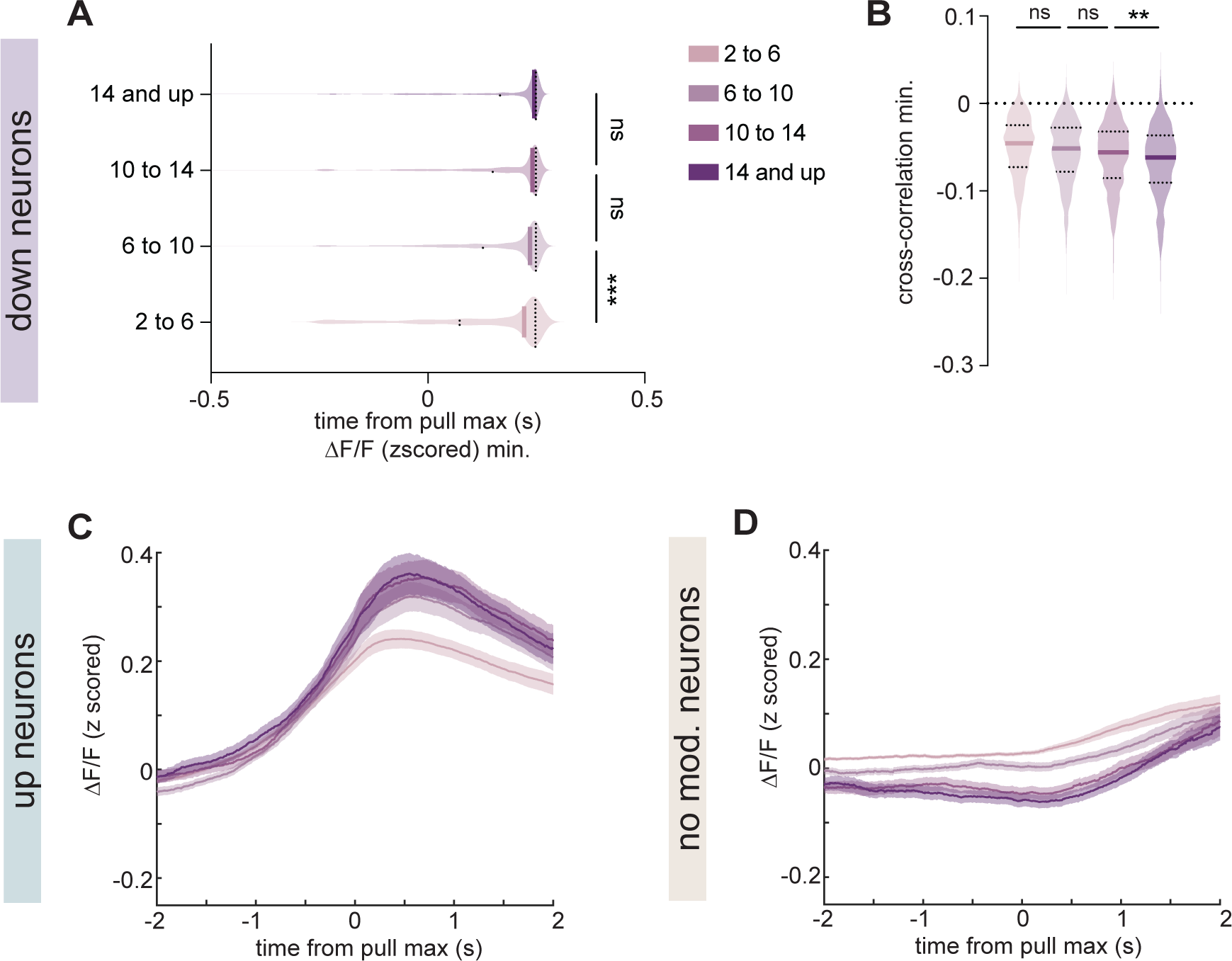
Features of M1^CT^ activity during pulls of varying velocity. As defined by ROC classification, units with decreases **(A,B)**, increases, n=168 **(C)**, or no modulation, n=168 **(D)** in activity during wheel pulls; legend for all as in **A**. **A,B** Distribution of time of ΔF/F minimum **(A)** or minimum of cross-correlation of wheel velocity to z-scored ΔF/F **(B)** during pulls of increasing velocity; thick, line, mean; thin lines, quartiles; One way ANOVA, multiple comparisons, p<0.0001, 2-6 v. 6 to 10; p=0.1759, 6-10 v. 10-14; p=0.8196, 10-14 v. 14 and up **(A)**. One way ANOVA, multiple comparisons, p=0.0711, 2-6 v. 6 to 10; p=0.0657, 6-10 v. 10-14; p=0.0.039, 10-14 v. 14 and up **(B)**. See Table 6 for p-values for all other comparisons. **C,D** Z-scored ΔF/F during pulls of increasing velocity; solid line, mean; shaded area, SEM.

**Figure S6.**
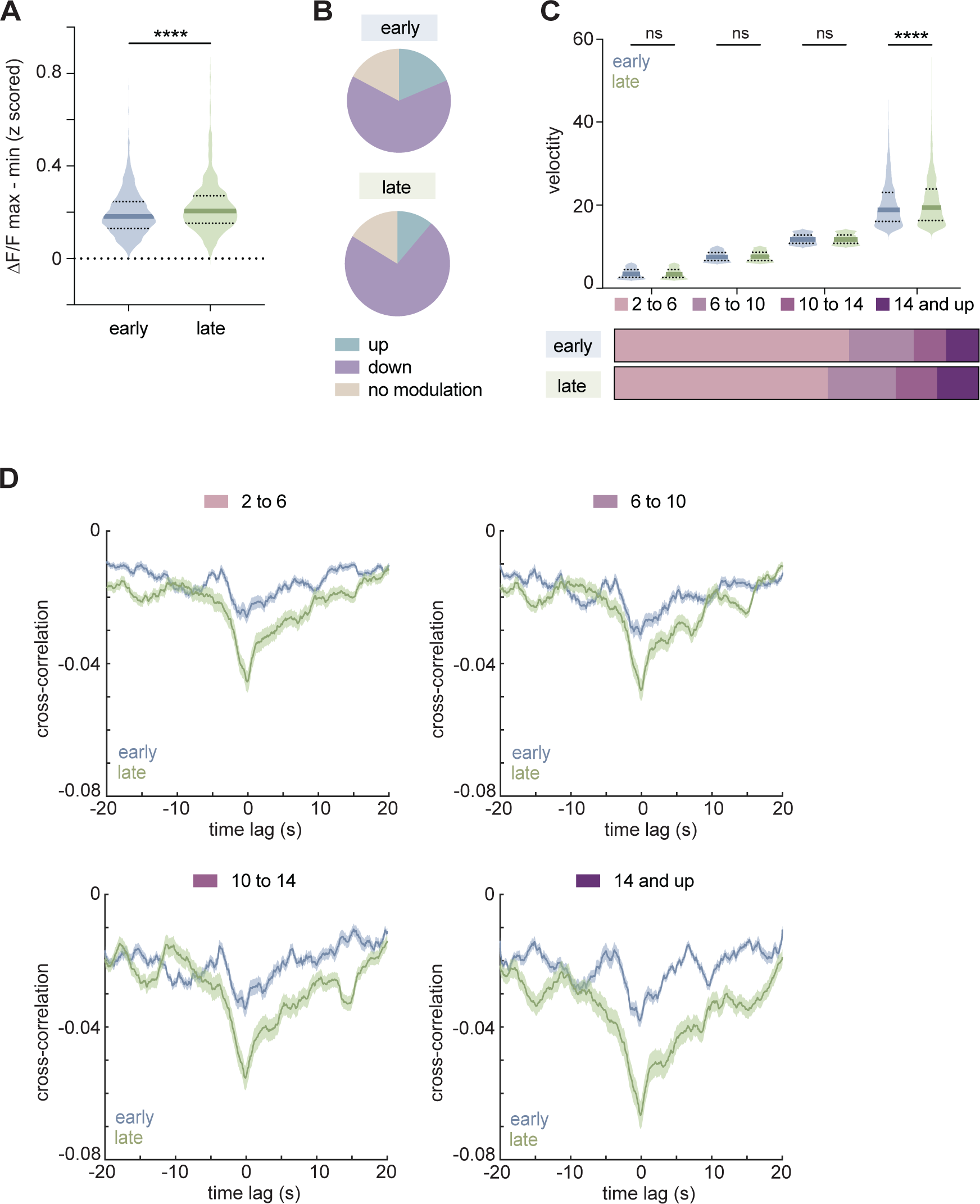
Comparison of M1^CT^ activity during early and late training. **A** Distribution of the difference in z-scored ΔF/F maximum and minimum before and during wheel pulls in down modulated neurons at early and late sessions; t-test, p<0.0001. **B** ROC based classification of all units over early and late sessions based on activity during wheel pulls. For early, n=237 down modulated units, n=69 up modulated units, n=64 no modulation units; for late, n=225 down modulated units, n=35 up modulated units, n=50 no modulation units. **C** Top, distribution of peak velocities in pulls of each velocity category in early (blue) or late (green) sessions; thick line, mean; thin lines, quartiles. One-way ANOVA, multiple comparisons, for early v. late, p=9890, 2-6; p>0.9999, 6-10 and 10-14; p<0.0001, 14 and up; see Table 7 for p-values for all other comparisons; Bottom, proportion of pulls in early or late sessions in each velocity category. **D** Cross-correlation of wheel pulls for early (blue) or late (green) training sessions of down modulated neurons to encoder velocity, grouped by increasing pull velocity; solid line, mean; shaded area, SEM. For **A-D**, early: n=370 units, late: n= 310 units from N=4 mice.

**Figure S7.**
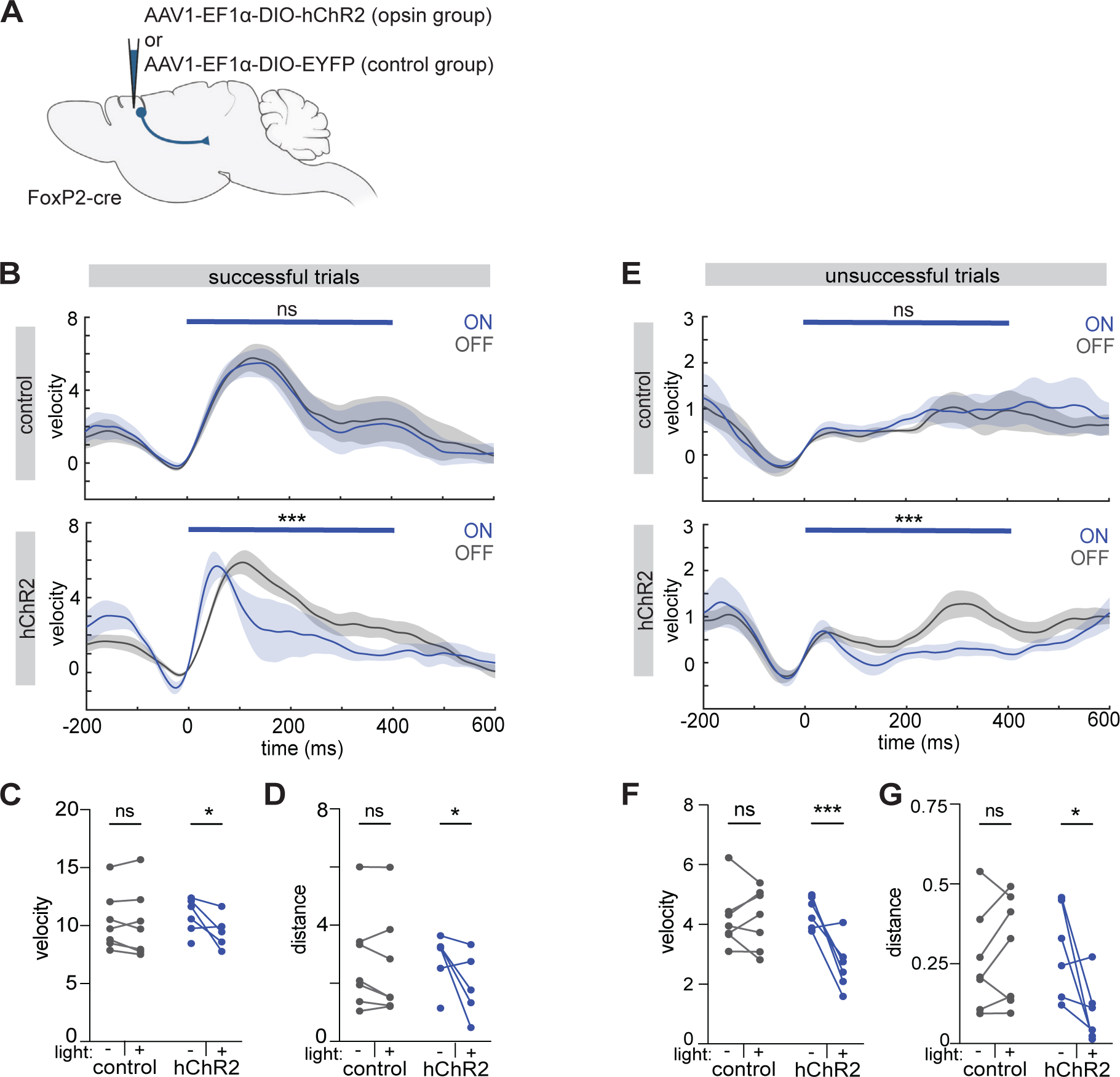
Closed-loop optogenetic perturbation at trial initiation. **A** Schematic of injection strategy for optogenetic experiments targeting M1^CT^ neurons **B,E** Velocity traces of successful **(B)** and unsuccessful **(E)** trials from control, top, and hChR2, bottom, animals during light, blue, and no light trials, gray, of closed-loop with light delivery at trial start; 0=trial start; blue bar denotes time of light; p>0.9999 (control), p<0.0001 (hChR2) **(B)**, p>0.9999 (control), p<0.0001 (hChR2) **(E)**, time x condition, two-way ANOVA; solid line, mean; shaded area, SEM. **C,F** Max velocity of successful trials **(C)** and unsuccessful trials **(F)** for each animal with light on or off in each group; two way ANOVA, multiple comparisons. **C**, p=0.9741 (control), p=0.0204 (hChR2)**; F**, p=0.9997 (control), p=0.0073 (hChR2); **D,G** Wheel distance traveled during successful **(D)** and unsuccessful **(G)** trials for each animal with light on or off for each group, **D,** Mixed-effect analysis, multiple comparisons, p=0.8655 (control), p=0.0163 (hChR2)**; G,** two way ANOVA, multiple comparisons; p=0.7514 (control), p=0.0145 (hChR2); N=7, control group; N=6, hChR2 group for **B-G.**

**Figure S8.**
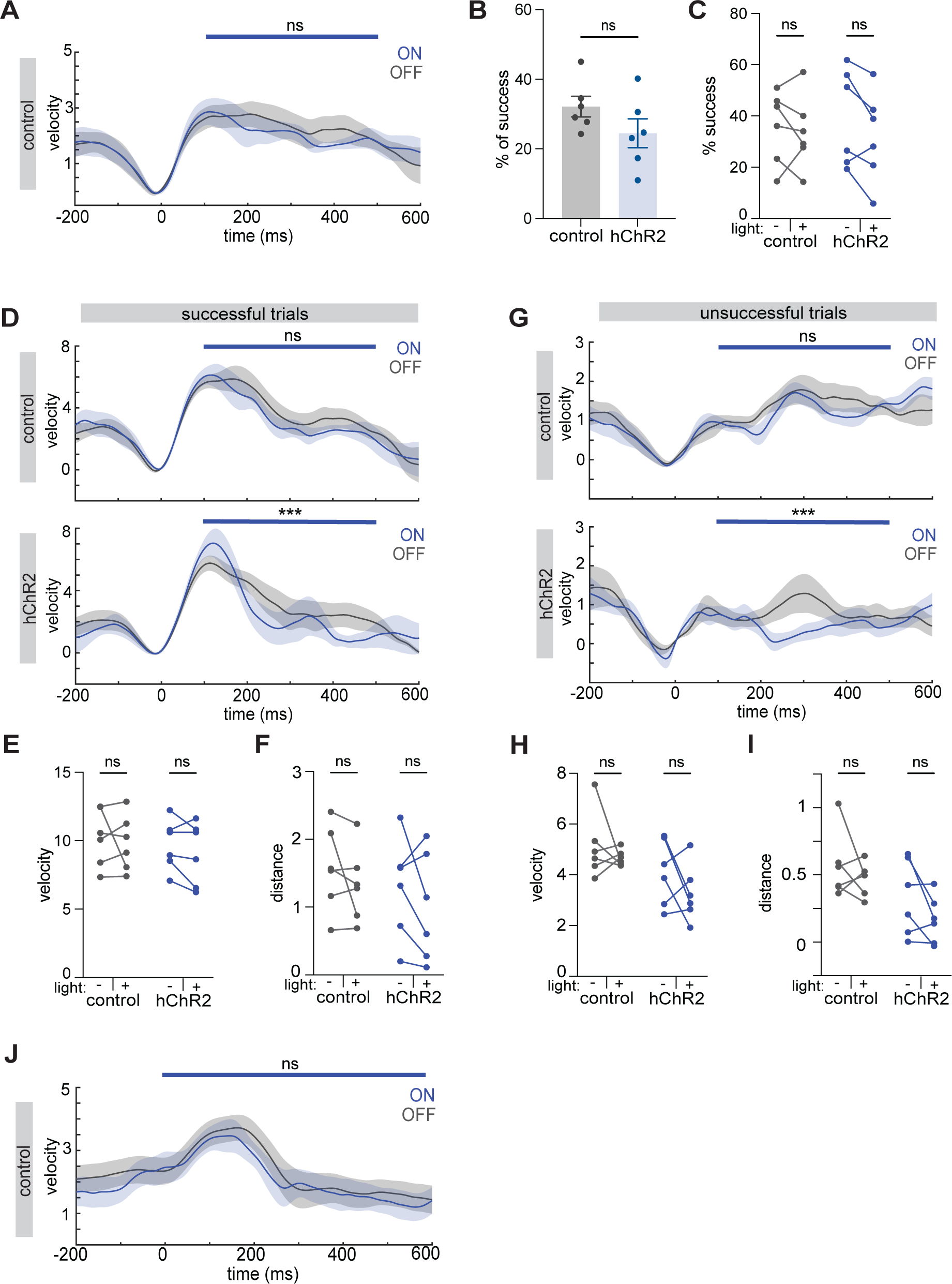
Optogenetic manipulations of ongoing trials or prior to trial start. **A** Velocity traces of control animals during light, blue, and no light trials, gray, of closed-loop with light delivery at trial maximum; 0=trial start; blue bar denotes time of light; p=0.8618, time x condition, two way ANOVA solid line, mean; shaded area, SEM. **B** Percent of successful trials with light on, (# successful trials during light on / # of all light on trials); control, gray; hChR2, blue; error bars, SEM; two-tailed unpaired t-test, p=0.1644. **C** Percentage of trials that are successful for each animal with light on or off in each group, (# (un)successful trials during light on / # of all (un)successful trials); two-way ANOVA, multiple comparisons; p=0.8417, control; p=0.1467, hChR2**. D,G** Velocity traces of successful **(D)** and unsuccessful **(G)** trials from control, top, and hChR2, bottom, animals during light, blue, and no light trials, gray, of closed-loop light delivery to max of trial; 0=trial start; blue bar denotes time of light; p=0.4331 (control), p<0.0001 (hChR2) **(D)**, p>0.9999 (control), p<0.0001 (hChR2) **(G)**, time x condition, two-way ANOVA solid line, mean; shaded area, SEM. **E,H** Max velocity of successful trials **(E)** and unsuccessful trials **(H)** for each animal with light on or off in each group; two way ANOVA, multiple comparisons. **E**, p=0.9835 (control), p=0.7817 (hChR2)**; H**, p=0.5749 (control), p=0.9991 (hChR2); **F,I** Wheel distance traveled during successful **(F)** and unsuccessful **(I)** trials for each animal with light on or off for each group; error bars, SEM; two way ANOVA, multiple comparisons, **F,** p=0.5232, (control), p=0.3938 (hChR2)**; I,** p=0.5969 (control), p=0.2309 (hChR2); **J** Velocity traces of all pulls from control animals during light, blue, and no light trials, gray, of open loop light delivery with 100 ms light preceding trial start; 0=trial start; blue bar denotes time of light; p>0.9999, time x condition, two way ANOVA solid line, mean; shaded area, SEM; N=11, each group. **A-I**: N=6, control group; N=6, hChR2 group.

**Figure S9.**
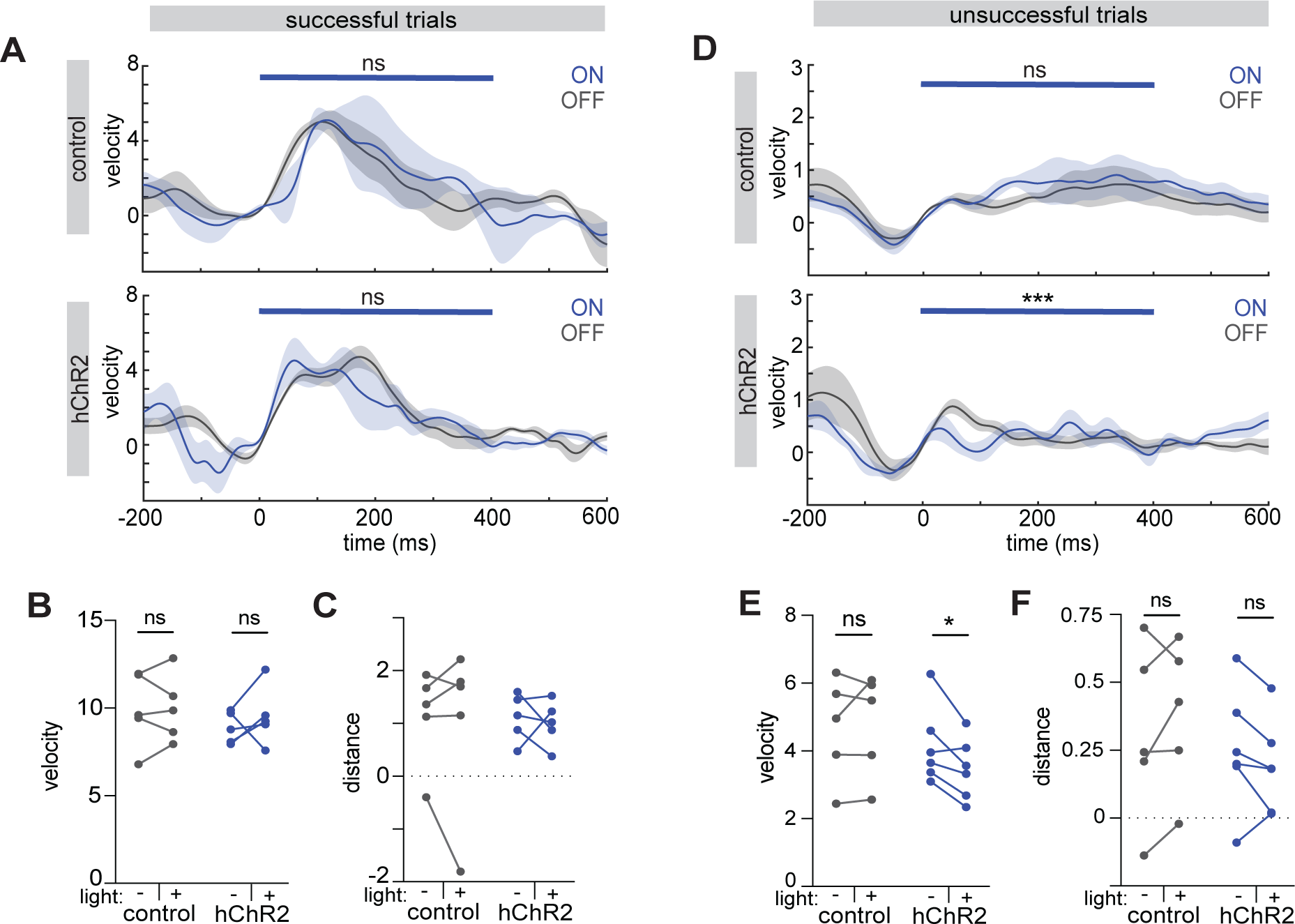
Optogenetic manipulations during early training. **A,D** Velocity traces of successful **(A)** and unsuccessful **(D)** trials from control, top, and hChR2, bottom, animals during light, blue, and no light trials, gray, of closed-loop with light delivery at trial start; 0=trial start; blue bar denotes time of light; p>0.9999 (control), p>0.9999 (hChR2) **(A)**, p>0.9999 (control), p<0.0001 (hChR2) **(D)**, time x condition, two-way ANOVA; solid line, mean; shaded area, SEM. **B,E** Max velocity of successful trials **(B)** and unsuccessful trials **(E)** for each animal with light on or off in each group; two way ANOVA, multiple comparisons, **B**, p=0.9958 (control), p=0.5515 (hChR2)**; E**, p=0.8443, (control), p=0.0316 (hChR2); **C,F** Wheel distance traveled during successful **(C)** and unsuccessful **(F)** trials for each animal with light on or off for each group; two way ANOVA, multiple comparisons, **C,** p=0.903, (control), p=0.9326, hChR2**; F,** p=0.3759 (control), p=0.3927 (hChR2); N=5, control group; N=5, hChR2 group for **A-C.** N=5, control group; N=6, hChR2 group for **D-F.**

**Figure S10.**
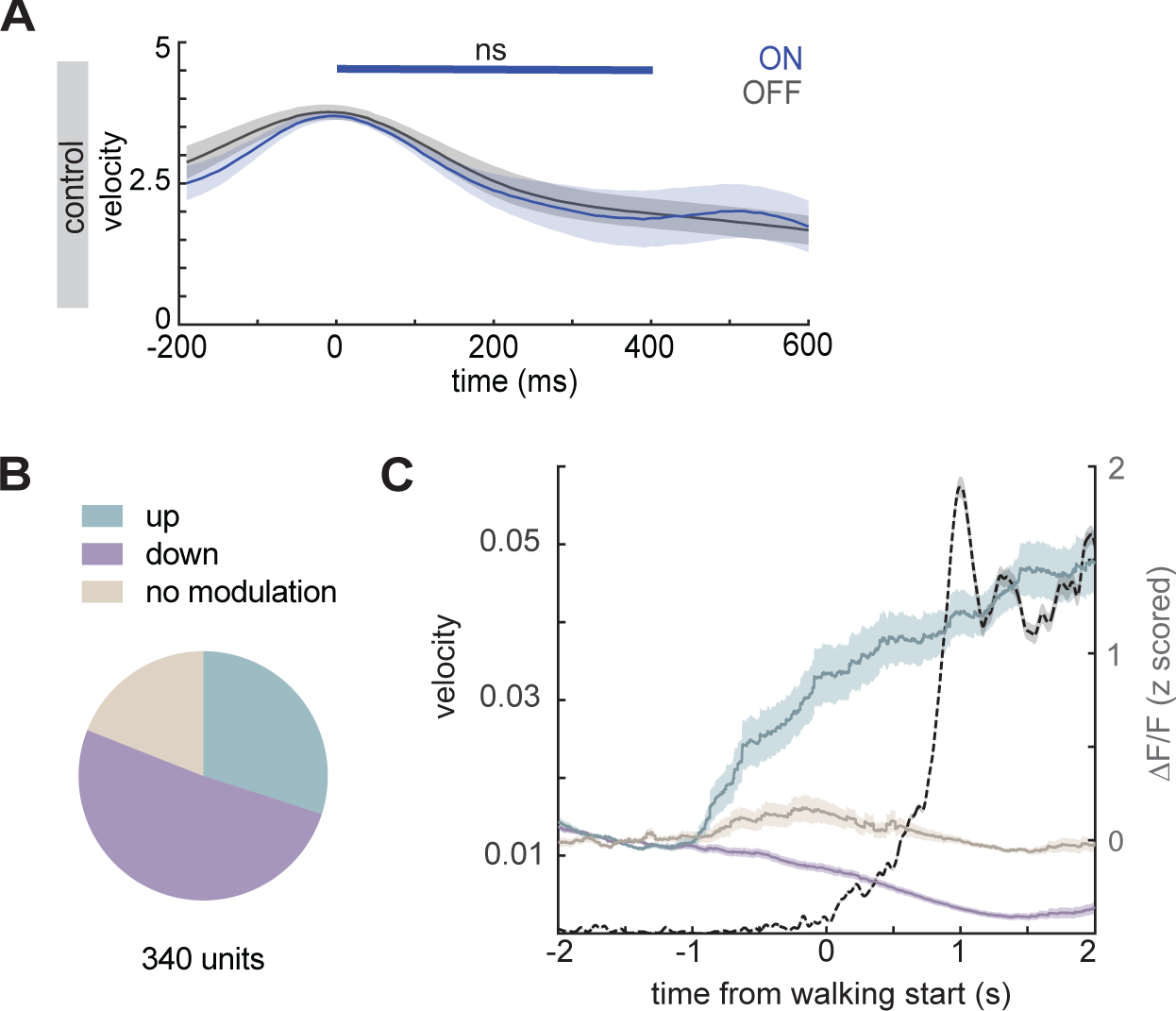
Optogenetic manipulations during walking. **A** Velocity traces of walking trials from control animals (see Figure 4H for hChR2 animals) during light, blue, and no light trials, gray; 0=trial start; blue bar denotes time of light, p>0.9999 (control), p=0.9958 (hChR2), time x condition, two way ANOVA; solid line, mean; shaded area, SEM. **B** ROC based classification of all units across sessions based on activity during walking bouts. **C** Z-scored ΔF/F of each group during walking bouts; solid lines, mean; shaded areas, SEM; wheel velocity, dotted line; 0=start of walking bout. For **B,C,** n=173 downmodulated units, n=102 upmodulated units, n=65 no modulation units; N=3 animals.

**Figure S11.**
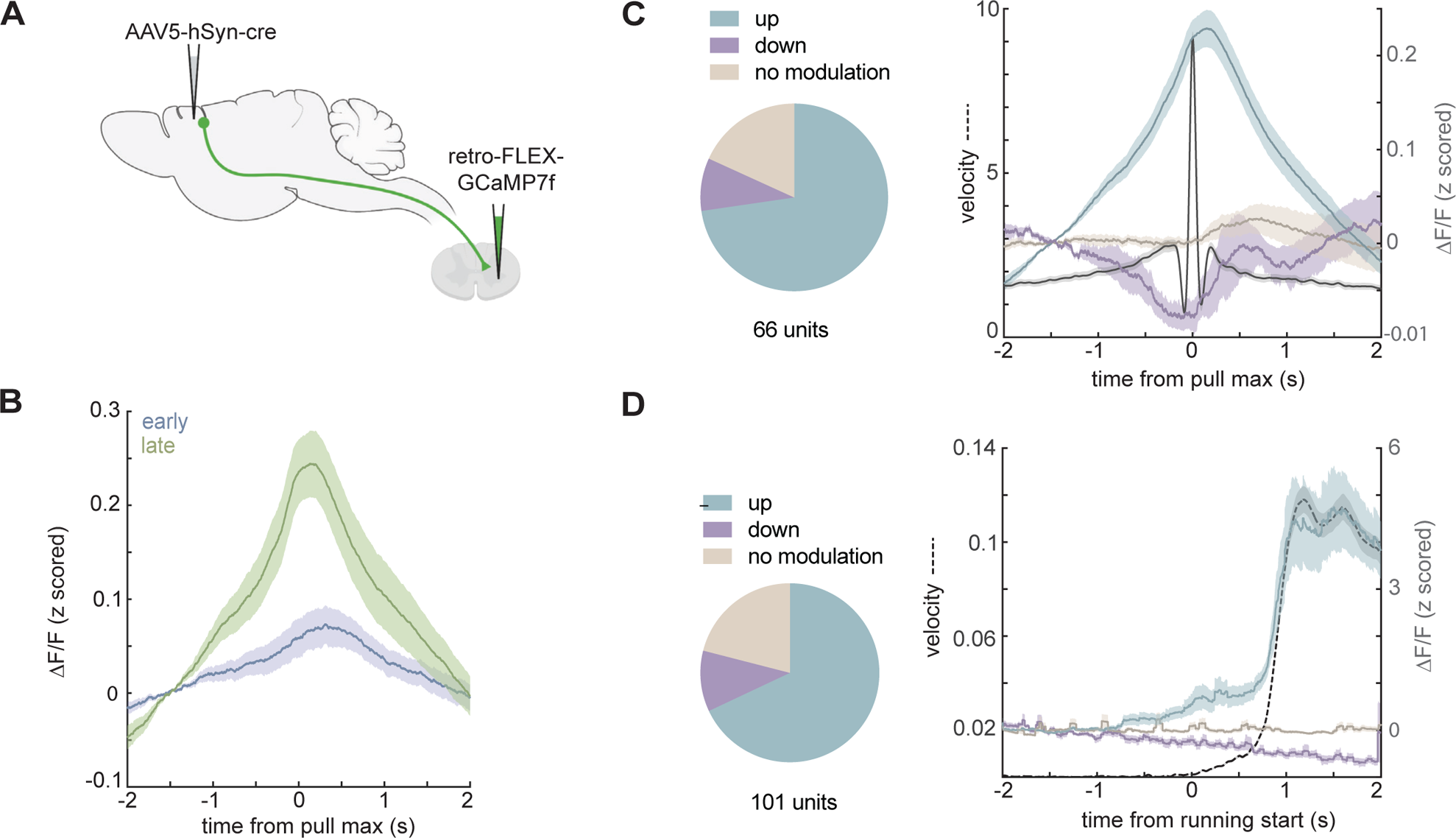
Calcium imaging of M1 corticospinal activity. **A** Schematic of injection strategy for calcium imaging of M1 corticospinal neurons. **B** Z-scored ΔF/F during wheel pulls at early, blue, or late, green, training, n=66 units, N=3 animals. **C,D.** Left, ROC based classification of all units across sessions based on activity during wheel pulls, n=48 upmodulated units, n=6 downmodulated units, n=12 no modulation units. **(C)** or during locomotion, n=74 upmodulated units, n=12 downmodulated units, n=23 no modulation units **(D)**. Right, Z-scored ΔF/F of each ROC group; solid lines, mean; shaded areas, SEM; wheel velocity during pulls, dotted line; 0=pull velocity max.

**Figure S12.**
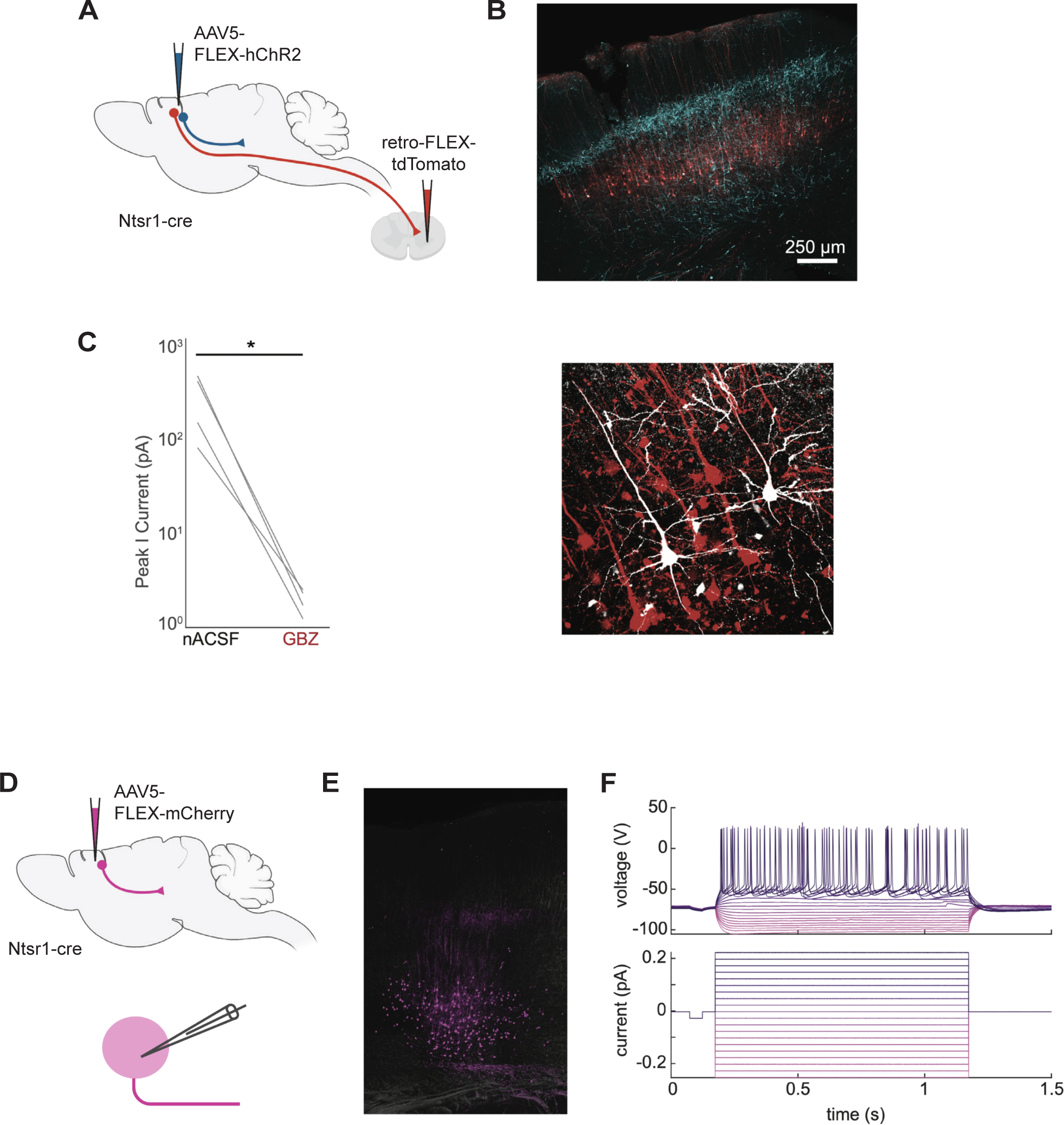
Slice electrophysiology of M1 corticospinal neurons during M1^CT^ photostimulation. **A** Schematic of injection strategy for labeling M1 corticospinal neurons and expressing channelrhodopsin (hChR2) in M1^CT^ neurons in Ntsr1-cre mice. **B** Expression of hChR2 (cyan) in M1^CT^ neurons and tdTomato in corticospinal neurons (red), top; Neurobiotin filled corticospinals post recording (white). **C** Peak inhibitory current with control treatment (nACSF) or GABAzine (GBZ); p=0.0284, paired t-test. **D** Schematic of injection strategy for labeling M1^CT^ neurons in Ntsr1-cre mice, top, for recordings of baseline firing properties, bottom. **E** Expression of mCherry in M1^CT^ neurons. **F** Firing of exemplar M1^CT^ neuron, top, at varying currents, bottom.

## REFERENCES

1 Arber, S. Motor circuits in action: specification, connectivity, and function. Neuron 74, 975–989 (2012). 10.1016/j.neuron.2012.05.011

2 Svoboda, K. & Li, N. Neural mechanisms of movement planning: motor cortex and beyond. Curr Opin Neurobiol 49, 33–41 (2018). 10.1016/j.conb.2017.10.023

3 Arber, S. & Costa, R. M. Connecting neuronal circuits for movement. Science 360, 1403–1404 (2018). 10.1126/science.aat5994

4 Jeong, M. et al. Comparative three-dimensional connectome map of motor cortical projections in the mouse brain. Sci Rep 6, 20072 (2016). 10.1038/srep20072

5 Whishaw, I. Q., Pellis, S. M., Gorny, B., Kolb, B. & Tetzlaff, W. Proximal and distal impairments in rat forelimb use in reaching follow unilateral pyramidal tract lesions. Behav Brain Res 56, 59–76 (1993). 10.1016/0166-4328(93)90022-i

6 Whishaw, I. Q., Pellis, S. M., Gorny, B. P. & Pellis, V. C. The impairments in reaching and the movements of compensation in rats with motor cortex lesions: an endpoint, videorecording, and movement notation analysis. Behav Brain Res 42, 77–91 (1991). 10.1016/s0166-4328(05)80042-7

7 Lawrence, D. G. & Kuypers, H. G. The functional organization of the motor system in the monkey. I. The effects of bilateral pyramidal lesions. Brain 91, 1–14 (1968). 10.1093/brain/91.1.1

8 Evarts, E. V. Pyramidal tract activity associated with a conditioned hand movement in the monkey. J Neurophysiol 29, 1011–1027 (1966). 10.1152/jn.1966.29.6.1011

9 Karni, A. et al. Functional MRI evidence for adult motor cortex plasticity during motor skill learning. Nature 377, 155–158 (1995). 10.1038/377155a0

10 Matsuzaka, Y., Picard, N. & Strick, P. L. Skill representation in the primary motor cortex after long-term practice. J Neurophysiol 97, 1819–1832 (2007). 10.1152/jn.00784.2006

11 Kawai, R. et al. Motor cortex is required for learning but not for executing a motor skill. Neuron 86, 800–812 (2015). 10.1016/j.neuron.2015.03.024

12 Miri, A. et al. Behaviorally Selective Engagement of Short-Latency Effector Pathways by Motor Cortex. Neuron 95, 683–696 e611 (2017). 10.1016/j.neuron.2017.06.042

13 Yang, G., Pan, F. & Gan, W. B. Stably maintained dendritic spines are associated with lifelong memories. Nature 462, 920–924 (2009). 10.1038/nature08577

14 Fu, M., Yu, X., Lu, J. & Zuo, Y. Repetitive motor learning induces coordinated formation of clustered dendritic spines in vivo. Nature 483, 92–95 (2012). 10.1038/nature10844

15 Peters, A. J., Chen, S. X. & Komiyama, T. Emergence of reproducible spatiotemporal activity during motor learning. Nature 510, 263–267 (2014). 10.1038/nature13235

16 Xu, T. et al. Rapid formation and selective stabilization of synapses for enduring motor memories. Nature (2009). 10.1038/nature08389

17 Harms, K. J., Rioult-Pedotti, M. S., Carter, D. R. & Dunaevsky, A. Transient spine expansion and learning-induced plasticity in layer 1 primary motor cortex. J Neurosci 28, 5686–5690 (2008). 10.1523/JNEUROSCI.0584-08.2008

18 Greenough, W. T., Larson, J. R. & Withers, G. S. Effects of unilateral and bilateral training in a reaching task on dendritic branching of neurons in the rat motor-sensory forelimb cortex. Behav Neural Biol 44, 301–314 (1985).

19 Withers, G. S. & Greenough, W. T. Reach training selectively alters dendritic branching in subpopulations of layer II-III pyramids in rat motor-somatosensory forelimb cortex. Neuropsychologia 27, 61–69 (1989). 10.1016/0028-3932(89)90090-0

20 Gloor, C., Luft, A. R. & Hosp, J. A. Biphasic plasticity of dendritic fields in layer V motor neurons in response to motor learning. Neurobiology of learning and memory (2015).

21 Wang, L., Conner, J. M., Rickert, J. & Tuszynski, M. H. Structural plasticity within highly specific neuronal populations identifies a unique parcellation of motor learning in the adult brain. Proc Natl Acad Sci U S A 108, 2545–2550 (2011). 10.1073/pnas.1014335108

22 Chen, S. X., Kim, A. N., Peters, A. J. & Komiyama, T. Subtype-specific plasticity of inhibitory circuits in motor cortex during motor learning. Nature neuroscience (2015). 10.1038/nn.4049

23 Peters, A. J., Lee, J., Hedrick, N. G., O’Neil, K. & Komiyama, T. Reorganization of corticospinal output during motor learning. Nat Neurosci 20, 1133–1141 (2017). 10.1038/nn.4596

24 Laubach, M., Wessberg, J. & Nicolelis, M. A. Cortical ensemble activity increasingly predicts behaviour outcomes during learning of a motor task. Nature 405, 567–571 (2000). 10.1038/35014604

25 Huber, D. et al. Multiple dynamic representations in the motor cortex during sensorimotor learning. Nature 484, 473–478 (2012). 10.1038/nature11039

26 Masamizu, Y. et al. Two distinct layer-specific dynamics of cortical ensembles during learning of a motor task. Nat Neurosci 17, 987–994 (2014). 10.1038/nn.3739

27 Nelson, A., Abdelmesih, B. & Costa, R. M. Corticospinal populations broadcast complex motor signals to coordinated spinal and striatal circuits. Nat Neurosci 24, 1721–1732 (2021). 10.1038/s41593-021-00939-w

28 Yao, Z. et al. A transcriptomic and epigenomic cell atlas of the mouse primary motor cortex. Nature 598, 103–110 (2021). 10.1038/s41586-021-03500-8

29 Network, B. I. C. C. A multimodal cell census and atlas of the mammalian primary motor cortex. Nature 598, 86–102 (2021). 10.1038/s41586-021-03950-0

30 Bakken, T. E. et al. Comparative cellular analysis of motor cortex in human, marmoset and mouse. Nature 598, 111–119 (2021). 10.1038/s41586-021-03465-8

31 Park, J. et al. Motor cortical output for skilled forelimb movement is selectively distributed across projection neuron classes. Sci Adv 8, eabj5167 (2022). 10.1126/sciadv.abj5167

32 Warriner, C. L., Fageiry, S., Saxena, S., Costa, R. M. & Miri, A. Motor cortical influence relies on task-specific activity covariation. bioRxiv, 2022.2002.2009.479479 (2022). 10.1101/2022.02.09.479479

33 Fosque, B. F. et al. Neural circuits. Labeling of active neural circuits in vivo with designed calcium integrators. Science (New York, N.Y.) 347, 755–760 (2015). 10.1126/science.1260922

34 DeNardo, L. A. et al. Temporal evolution of cortical ensembles promoting remote memory retrieval. Nat Neurosci 22, 460–469 (2019). 10.1038/s41593-018-0318-7

35 Wang, W. et al. A light- and calcium-gated transcription factor for imaging and manipulating activated neurons. Nat Biotechnol (2017). 10.1038/nbt.3909

36 Lee, D., Hyun, J. H., Jung, K., Hannan, P. & Kwon, H. B. A calcium- and light-gated switch to induce gene expression in activated neurons. Nature Biotechnology 35, 858-+ (2017). 10.1038/nbt.3902

37 Kim, C. K. et al. A Molecular Calcium Integrator Reveals a Striatal Cell Type Driving Aversion. Cell 183, 2003–2019 e2016 (2020). 10.1016/j.cell.2020.11.015

38 Tennant, K. A. et al. The organization of the forelimb representation of the C57BL/6 mouse motor cortex as defined by intracortical microstimulation and cytoarchitecture. Cereb Cortex 21, 865–876 (2011). 10.1093/cercor/bhq159

39 Hira, R. et al. In vivo optogenetic tracing of functional corticocortical connections between motor forelimb areas. Front Neural Circuits 7, 55 (2013). 10.3389/fncir.2013.00055

40 Yamawaki, N. & Shepherd, G. M. Synaptic circuit organization of motor corticothalamic neurons. J Neurosci 35, 2293–2307 (2015). 10.1523/JNEUROSCI.4023-14.2015

41 Munoz-Castaneda, R. et al. Cellular anatomy of the mouse primary motor cortex. Nature 598, 159–166 (2021). 10.1038/s41586-021-03970-w

42 Harris, J. A. et al. Hierarchical organization of cortical and thalamic connectivity. Nature 575, 195–202 (2019). 10.1038/s41586-019-1716-z

43 Canavan, A. G., Nixon, P. D. & Passingham, R. E. Motor learning in monkeys (Macaca fascicularis) with lesions in motor thalamus. Exp Brain Res 77, 113–126 (1989). 10.1007/BF00250573

44 Bosch-Bouju, C., Hyland, B. I. & Parr-Brownlie, L. C. Motor thalamus integration of cortical, cerebellar and basal ganglia information: implications for normal and parkinsonian conditions. Front Comput Neurosci 7, 163 (2013). 10.3389/fncom.2013.00163

45 Johnson, F. & Bottjer, S. W. Induced cell death in a thalamic nucleus during a restricted period of zebra finch vocal development. J Neurosci 13, 2452–2462 (1993).

46 Sauerbrei, B. A. et al. Cortical pattern generation during dexterous movement is input-driven. Nature 577, 386–391 (2020). 10.1038/s41586-019-1869-9

47 Tanaka, Y. H. et al. Thalamocortical Axonal Activity in Motor Cortex Exhibits Layer-Specific Dynamics during Motor Learning. Neuron 100, 244–258 e212 (2018). 10.1016/j.neuron.2018.08.016

48 Pnevmatikakis, E. A. et al. Simultaneous Denoising, Deconvolution, and Demixing of Calcium Imaging Data. Neuron 89, 285–299 (2016). 10.1016/j.neuron.2015.11.037

49 Mittmann, W. et al. Two-photon calcium imaging of evoked activity from L5 somatosensory neurons in vivo. Nat Neurosci 14, 1089–1093 (2011). 10.1038/nn.2879

50 Arber, S. & Costa, R. M. Networking brainstem and basal ganglia circuits for movement. Nat Rev Neurosci 23, 342–360 (2022). 10.1038/s41583-022-00581-w

51 Bortone, D. S., Olsen, S. R. & Scanziani, M. Translaminar inhibitory cells recruited by layer 6 corticothalamic neurons suppress visual cortex. Neuron 82, 474–485 (2014). 10.1016/j.neuron.2014.02.021

52 Olsen, S. R., Bortone, D. S., Adesnik, H. & Scanziani, M. Gain control by layer six in cortical circuits of vision. Nature 483, 47–52 (2012). 10.1038/nature10835

53 Pauzin, F. P. & Krieger, P. A Corticothalamic Circuit for Refining Tactile Encoding. Cell Rep 23, 1314–1325 (2018). 10.1016/j.celrep.2018.03.128

54 Ziegler, K. et al. Primary somatosensory cortex bidirectionally modulates sensory gain and nociceptive behavior in a layer-specific manner. Nat Commun 14, 2999 (2023). 10.1038/s41467-023-38798-7

55 Hrvatin, S. et al. Single-cell analysis of experience-dependent transcriptomic states in the mouse visual cortex. Nat Neurosci 21, 120–129 (2018). 10.1038/s41593-017-0029-5

56 Chi, J., Crane, A., Wu, Z. & Cohen, P. Adipo-Clear: A Tissue Clearing Method for Three-Dimensional Imaging of Adipose Tissue. J Vis Exp (2018). 10.3791/58271

57 Renier, N. et al. Mapping of Brain Activity by Automated Volume Analysis of Immediate Early Genes. Cell 165, 1789–1802 (2016). 10.1016/j.cell.2016.05.007

58 Botta, P. et al. An Amygdala Circuit Mediates Experience-Dependent Momentary Arrests during Exploration. Cell 183, 605–619 e622 (2020). 10.1016/j.cell.2020.09.023

