## supplemental tables for "Corticothalamic Neurons in Motor Cortex Have a Permissive Role in Motor Execution"

**Table 1**

| summary metrics | |
|---|---|
| median UMI count | 7.97E+03 |
| median gene count | 3503 |
| total cell count | 16490 |
| # runs | |
| control | 4 |
| early training | 4 |
| late training | 5 |

| | control | early | late |
|---|---|---|---|
| median UMI count | 8557 | 6198 | 10390 |
| median gene count | 3694 | 2911 | 4035 |
| total cell count | 5369 | 5493 | 2923 |

**Table 2**

| Figure 1G | | | | | | | | | | | | |
|---|---|---|---|---|---|---|---|---|---|---|---|---|
| | ANCOM-BC | | | | | | MASC | | | | | |
| | pvalue_early_vs_control | pvalue_late_vs_control | pvalue_late_vs_early | logfc_early_vs_control | logfc_late_vs_control | logfc_late_vs_early | pvalue_early_vs_control | pvalue_late_vs_control | pvalue_late_vs_early | oddsratio_early_vs_control | oddsratio_late_vs_control | oddsratio_late_vs_early |
| L2/3T_Cux2+_ | 0.314311545 | 0.688105652 | 0.253692781 | -0.27004054 | -0.087459919 | -0.318023972 | 0.416326 | 0.32622428 | 0.800263109 | 0.833414037 | 0.780734882 | 0.937181992 |
| L5/6IT_Fezf2+_Ccdc80+ | 0.385351853 | 0.00037104 | 0.573116669 | -0.126061349 | 0.444062957 | -0.188350382 | 0.740895886 | 0.00976976 | 0.010411388 | 0.949588549 | 1.416934229 | 1.520025651 |
| L5/6IT_Rorb+_Adam33+_Hspb3+ | 0.90014996 | 0.597398862 | 0.005823879 | 0.027043827 | -0.163327305 | 0.069519713 | 0.58419304 | 0.39028402 | 0.147404364 | 1.182348225 | 0.7390475 | 0.635826562 |
| L5/6IT_Rorb+_Atp5o+ | 0.145123315 | 0.008802587 | 0.705605536 | 1.266158392 | 1.439437956 | -0.690975725 | 0.116900365 | 0.03772686 | 0.84730077 | 6.134906486 | 4.984313327 | 0.828033627 |
| L5/6NP_Fezf2+_Tshz2+ | 0.425587444 | 0.105252944 | 0.100516587 | 0.24461935 | 0.217618573 | -0.327325029 | 0.232518644 | 0.34854178 | 0.32670228 | 1.560564324 | 1.17744817 | 0.70586913 |
| L5ET_Fezf2+_Bcl6+ | 0.383289507 | 0.00734846 | 0.705718194 | 0.134888619 | 0.557043029 | -0.52760537 | 0.210389024 | 0.1178491 | 0.601390916 | 1.26967093 | 1.420827341 | 1.126579075 |
| L5IT_Rorb+_Adam33+_Trpm4+ | 0.19140785 | 0.680489443 | 0.60115558 | -0.403049609 | -0.091195299 | -0.078450183 | 0.348991033 | 0.41204097 | 0.807554049 | 0.727333927 | 0.82580768 | 1.093021211 |
| L5IT_Rorb+_Atp6v0c+ | 0.028427845 | 0.493241946 | 0.489743906 | -0.486595739 | -0.174341528 | -0.188750283 | 0.060755413 | 0.32783415 | 0.755340008 | 0.630843701 | 0.698948444 | 1.115822996 |
| L6CT_Fezf2+_Foxp2+ | 0.157494187 | 0.000110259 | 0.21600365 | 0.443666249 | 1.268605833 | 0.324334991 | 0.150257429 | 0.00606913 | 0.055825494 | 1.594822427 | 3.234999561 | 1.977640151 |
| Ndnf_Chst7 | 0.005723362 | 0.41307281 | 0.003943091 | -0.784863684 | 0.097239296 | 0.79986784 | 0.005646497 | 0.79614327 | 0.005335598 | 0.435618518 | 0.94488477 | 2.169064748 |
| Ndnf_Zfp804b | 0.21751502 | 0.002153044 | 0.144097542 | 0.47773345 | 1.071174499 | 0.51120591 | 0.131720134 | 0.03727113 | 0.438800562 | 2.927631186 | 4.543830429 | 1.580608086 |
| Pvalb | 0.837784536 | 0.501073753 | 0.573031189 | -0.048395344 | 0.197318547 | 0.163478752 | 0.646929224 | 0.92150994 | 0.589054265 | 0.742616034 | 1.064986931 | 1.422775883 |
| Sst_Gabrd | 3.92E-06 | 3.55E-09 | 0.765734322 | 0.628617206 | 0.760387669 | 0.049535323 | 0.103857051 | 0.06774033 | 0.899161316 | 3.461538455 | 3.711341145 | 1.072164966 |
| Sst_Htr1a | 0.373933136 | 0.640489206 | 0.672008462 | -0.462302168 | -0.136082159 | 0.243984869 | 0.647783734 | 0.81161573 | 0.943938043 | 0.68168327 | 0.902075959 | 1.071425779 |
| Sst_Myh8 | 0.212687719 | 0.56467028 | 0.355334223 | 0.455330411 | 0.231177058 | -0.306388492 | 0.449625522 | 0.98325938 | 0.192586698 | 1.824211149 | 0.972259802 | 0.3919598 |
| Vip_Pkib | 0.385883624 | 0.33111238 | 0.887593023 | 0.405162737 | 0.413688604 | -0.073709273 | 0.098235392 | 0.36349801 | 0.685390086 | 2.13889768 | 1.637975486 | 0.778707863 |
| Vip_Sema5b | 0.837252482 | 0.175984791 | 0.228396087 | -0.043240523 | -0.175417451 | -0.214412067 | 0.601515045 | 0.01574876 | 0.126761722 | 0.861016953 | 0.552481841 | 0.627402585 |
| Vip_Sncg | 0.152802578 | 0.020819684 | 0.489635731 | 0.48477617 | 0.702851254 | 0.135839945 | 0.087964822 | 0.07393333 | 0.863619894 | 4.064516129 | 3.711340206 | 0.913107511 |
| Vip_Syt9 | 0.693621108 | 6.20E-06 | 0.011416114 | -0.090870104 | -0.68169258 | -0.673057616 | 0.571950693 | 0.00378455 | 0.022005831 | 0.739316239 | 0.095632971 | 0.129353234 |

**Table 3**

| Figure 2I | |
|---|---|
| 2-6 v 6-10 | <0.0001 |
| 2-6 v 10-14 | <0.0001 |
| 2-6 v 14+ | <0.0001 |
| 6-10 v 10-14 | <0.0001 |
| 6-10 v 14+ | <0.0001 |
| 10-14 v 14+ | 0.0553 |

| Figure 2J | |
|---|---|
| 2-6 v 6-10 | <0.0001 |
| 2-6 v 10-14 | <0.0001 |
| 2-6 v 14+ | <0.0001 |
| 6-10 v 10-14 | 0.023 |
| 6-10 v 14+ | 0.0482 |
| 10-14 v 14+ | 0.9937 |

| Figure 2M | |
|---|---|
| 2-6 early v 2-6 late | <0.0001 |
| 2-6 early v 6-10 early | <0.0001 |
| 2-6 early v 6-10 late | <0.0001 |
| 2-6 early v 10-14 early | <0.0001 |
| 2-6 early v 10-14 late | <0.0001 |
| 2-6 early v 14+ early | <0.0001 |
| 2-6 early v. 14+ late | <0.0001 |
| 2-6_late v 6-10 early | 0.9994 |
| 2-6 late v 6-10 late | <0.0001 |
| 2-6 late v. 10=14 early | <0.0001 |
| 2-6 late v 10-14 late | <0.0001 |
| 2-6 late v 14+ early | <0.0001 |
| 2-6 late v 14+ late | <0.0001 |
| 6-10 early v 6-10 late | <0.0001 |
| 6-10 early v 10-14 early | <0.0001 |
| 6-10 early v 10-14 late | <0.0001 |
| 6-10 early v 14+ early | <0.0001 |
| 6-10 early v 14+ late | <0.0001 |
| 6-10 late v 10-14 early | 0.9994 |
| 6-10 late v 10-14 late | <0.0001 |
| 6-10 late v 14+ early | 0.005 |
| 6-10 late v 14+ late | <0.0001 |
| 10-14 early v 10-14 late | 0.0005 |
| 10-14 early v 14+ early | 0.0318 |
| 10-14 early v 14+ late | <0.0001 |
| 10-14 late v 14+ early | 0.9582 |
| 10-14 late v 14+ late | <0.0001 |
| 14+ early vs. 14+ late | <0.0001 |

**Table 4**

| Figure 5H | |
|---|---|
| untrained:excitatory vs. untrained:inhibitory | 0.9169 |
| untrained:excitatory vs. trained:excitatory | >0.9999 |
| untrained:excitatory vs. trained:inhibitory | <0.0001 |
| untrained:inhibitory vs. trained:excitatory | 0.9757 |
| untrained:inhibitory vs. trained:inhibitory | <0.0001 |
| trained:excitatory vs. trained:inhibitory | <0.0001 |

**Table 5**

| Figure S3F (right) | | |
|---|---|---|
| | LAYER V | LAYER VI |
| | anterior | |
| lateral v. ventral | <0.0001 | 0.0362 |
| lateral v medial | <0.0001 | 0.0212 |
| ventral v medial | 0.8207 | 0.9639 |
| | posterior | |
| lateral v. ventral | 0.5017 | 0.2431 |
| lateral v medial | 0.0173 | 0.6785 |
| ventral v medial | 0.0014 | 0.0533 |

**Table 6**

| Figure S5A | |
|---|---|
| 2-6 v 6-10 | 0.0004 |
| 2-6 v 10-14 | 0.0001 |
| 2-6 v 14+ | <0.0001 |
| 6-10 v 10-14 | 0.9962 |
| 6-10 v 14+ | 0.8323 |
| 10-14 v 14+ | 0.9241 |

| Figure S5B | |
|---|---|
| 2-6 v 6-10 | 0.0711 |
| 2-6 v 10-14 | <0.0001 |
| 2-6 v 14+ | <0.0001 |
| 6-10 v 10-14 | 0.0657 |
| 6-10 v 14+ | <0.0001 |
| 10-14 v 14+ | 0.0039 |

**Table 7**

| Figure S6C | |
|---|---|
| 2-6 early v 2-6 late | 0.989 |
| 2-6 early v 6-10 early | <0.0001 |
| 2-6 early v 6-10 late | <0.0001 |
| 2-6 early v 10-14 early | <0.0001 |
| 2-6 early v 10-14 late | <0.0001 |
| 2-6 early v 14+ early | <0.0001 |
| 2-6 early v. 14+ late | <0.0001 |
| 2-6_late v 6-10 early | <0.0001 |
| 2-6 late v 6-10 late | <0.0001 |
| 2-6 late v. 10=14 early | <0.0001 |
| 2-6 late v 10-14 late | <0.0001 |
| 2-6 late v 14+ early | <0.0001 |
| 2-6 late v 14+ late | <0.0001 |
| 6-10 early v 6-10 late | >0.9999 |
| 6-10 early v 10-14 early | <0.0001 |
| 6-10 early v 10-14 late | <0.0001 |
| 6-10 early v 14+ early | <0.0001 |
| 6-10 early v 14+ late | <0.0001 |
| 6-10 late v 10-14 early | <0.0001 |
| 6-10 late v 10-14 late | <0.0001 |
| 6-10 late v 14+ early | <0.0001 |
| 6-10 late v 14+ late | <0.0001 |
| 10-14 early v 10-14 late | >0.9999 |
| 10-14 early v 14+ early | <0.0001 |
| 10-14 early v 14+ late | <0.0001 |
| 10-14 late v 14+ early | <0.0001 |
| 10-14 late v 14+ late | <0.0001 |
| 14+ early vs. 14+ late | <0.0001 |
